# *In-situ* structures of the *Legionella* Dot/Icm T4SS identify the DotA-IcmX complex as the gatekeeper for effector translocation

**DOI:** 10.1101/2025.06.23.660953

**Authors:** Jian Yue, Samira Heydari, Donghyun Park, David Chetrit, Shoichi Tachiyama, Wangbiao Guo, Jack M. Botting, Shenping Wu, Craig R. Roy, Jun Liu

## Abstract

The Dot/Icm machine in *Legionella pneumophila* is one of the most versatile type IV secretion systems (T4SSs), with a remarkable capacity to translocate over 330 different effector proteins across the bacterial envelope into host cells. At least 27 Dot and Icm proteins are required for assembly and function of the system, yet the architecture and activation mechanism remain poorly understood at the molecular level. Here, we deploy cryo-electron microscopy to reveal *in-situ* structures of the Dot/Icm machine at near-atomic resolution. Importantly, two proteins essential for effector translocation, DotA and IcmX, form a pentameric protochannel at an inactive state. Upon activation, the DotA-IcmX protochannel undergoes extensive rearrangements to form an extended transenvelope passage capable of transporting effector proteins from the bacterial cytoplasm into host cells as revealed by cryo-electron tomography. Collectively, our findings suggest that the DotA-IcmX complex functions as the gatekeeper for effector translocation of the Dot/Icm T4SS.

## Introduction

Bacteria have evolved diverse, specialized secretion systems that transport macromolecules across the cell envelope(1, 2). Type IV secretion systems (T4SSs) are among the most versatile nanomachines, capable of translocating proteins and nucleoprotein complexes into a wide range of prokaryotic and eukaryotic recipient cells(3, 4). This versatility allows T4SSs to play critical roles in horizontal gene transfer, interbacterial competition(5–7), and pathogenesis, by delivering effectors into both prokaryotic and eukaryotic cells(8–10). Most DNA conjugation machines are categorized as type IVA secretion systems(11–15). Well-characterized examples include the *Agrobacterium tumefaciens* VirB/VirD4 system and the *Escherichia coli* R388 and pKM101 plasmid conjugation systems(16, 17). The Dot/Icm T4SS of *Legionella pneumophila* is a type IVB system markedly more complex and expanded(18–21). A hallmark of the Dot/Icm T4SS is its remarkable ability to translocate over 330 effector proteins – approximately 10% of the *L. pneumophila* proteome – into eukaryotic host cells(22, 23). These effectors manipulate host processes to establish a replication-permissive compartment, the *Legionella*-containing vacuole (LCV), essential for intracellular survival and the development of Legionnaires’ disease(24–27). At least 27 different Dot and Icm proteins are required for the assembly of the Dot/Icm machine and translocation of effector proteins into host cells(18).

Over the past two decades, structural and mechanistic studies have advanced our understanding of the T4SS superfamily’s architectural and functional diversity(3, 4, 28, 29). In particular, cryo-electron microscopy (cryo-EM) and cryo-electron tomography (cryo-ET) studies reveal that the Dot/Icm machine is a multi-layered complex with a characteristic “Wi-Fi” architecture(30) in the periplasmic space and unique cytoplasmic hexameric ATPase complexes(31). The outer membrane core complex (OMCC) exhibits three layers characterized by symmetry mismatches: a dome-shaped layer with 16-fold symmetry, a 13-fold symmetric outer membrane cap (OMC), and an 18-fold symmetric periplasmic ring (PR). The OMCC is composed of DotC, DotD, DotF, DotG (C-terminal), DotH, and DotK, along with three species-specific components, Dis1, Dis2, and Dis3(32, 33). Intriguingly, these previous studies have suggested that the OMCC is connected to the cytoplasmic ATPase complexes through a “plug-like” structure and a cylindrical periplasmic channel that spans the inner membrane(31, 34). This unique architecture suggested a pathway for the translocation of substrates through the Dot/Icm machine across the cell envelope(31, 34). However, low-resolution structural information of the cytoplasmic and periplasmic elements have prevented a complete mechanistic understanding of the assembly and function of the Dot/Icm machine at the molecular level.

In this study, we present the *L. pneumophila* Dot/Icm machine in a native cellular context at an unprecedented resolution by combining *in-situ* cryo-EM with cryo-ET and subtomogram averaging. Our near-atomic cryo-EM structure reveal that two proteins essential for effector translocation, DotA and IcmX, from a pentameric protochannel in the central axis of the intact Dot/Icm machine for the first time. Cryo-ET and subtomogram averaging further capture the activated state of the secretion machine, demonstrating that a major conformational change in the DotA-IcmX protochannel is required to form an extended transenvelope channel and to enable direct translocation of substrates from the bacterial cytoplasm into recipient cells. These findings offer fundamental insights into the architecture and activation mechanisms of T4SSs and establish a structural framework for more fully understanding T4SS-mediated host-pathogen interactions in *L. pneumophila* and many other bacteria.

## Results

### *In-situ* cryo-EM structure of the Dot/Icm machine reveals the DotA-IcmX complex

To gain a better understanding of the composition and assembly of the Dot/Icm machine, we employed an *in-situ* cryo-EM approach to determine the structure of Dot/Icm machines in frozen-hydrated bacterial cells (Fig. 1A). This approach preserved the structural integrity of the Dot/Icm machine within its native bacterial envelope, thereby avoiding disruption during traditional purification. Using a combined strategy of template matching(35) and deep learning(36) for particle selection, we identified and analyzed approximately 76,000 Dot/Icm machines from ∼49,000 images collected from *L. pneumophila* cell poles (SI Appendix, Fig. S1, S2; SI Appendix, Table S1). The final *in-situ* structures of the Dot/Icm machine were determined at resolutions ranging from ∼3.0 Å to 4.6 Å (Figs. 1A-1E; SI Appendix, Fig. S3).

**Figure 1.**
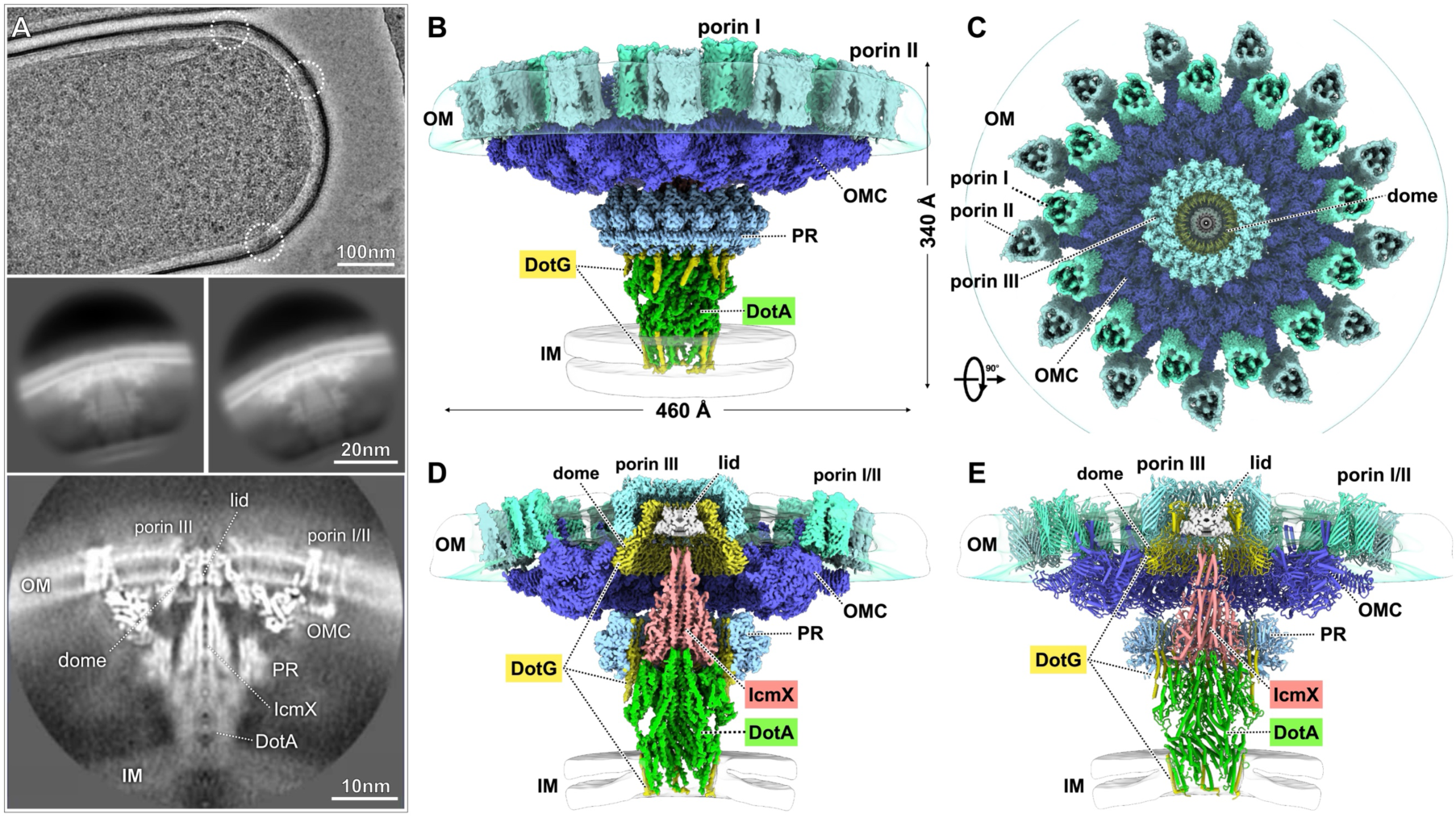
*In-situ* structure of the Dot/Icm machine at near-atomic resolution. (**A**) A representative cryo-EM micrograph shows the Dot/Icm machines localized at the bacterial cell pole, indicated by white circles. Representative 2D class averages and a central cross-section of the 3D average of Dot/Icm particles are displayed in the bottom panel. (**B**) Overall *in-situ* structure of the Dot/Icm machine. The structure spans 460 Å in width and 340 Å in height, extending across the bacterial envelope. Subcomplexes of outer membrane core complex (OMCC, including the outer membrane cap (OMC), periplasmic ring (PR), Dome), DotA, DotG and porins, are rendered in distinct colors. Outer membrane (OM) and inner membrane (IM) are shown as transparent layers. (**C**) Top view of the cryo-EM structure showing the OMCC and its association with porin arrays. Three porin arrays are integrated with the OMCC, anchoring it to the OM. Porin I and Porin II each form trimers with C13 symmetry at the OMC periphery and are shown in cyan and dark cyan, respectively. Porin III forms a ring with C16 symmetry, surrounding the dome. (**D**) Cross-sectional view of the cryo-EM map illustrating the structural organization of each subcomplex within the Dot/Icm machine. (**E**) Atomic model of the Dot/Icm machine.

The *in-situ* structures enabled us to build models for 13 known Dot/Icm proteins (Fig. 1E; SI Appendix, Fig. S4) as well as identify new components (Fig. 1E; SI Appendix, Table S2). The total complex resolved in our cryo-EM structures is approximately 8.5 MDa, 460 Å wide and 340 Å tall, spanning the entire bacterial envelope (Figs. 1B-1E). Importantly, our *in-situ* structures revealed for the first time three porin arrays associated with the OMCC in the outer membrane and the DotA-IcmX pentameric complex at the central axis of the Dot/Icm machine. The DotA-IcmX complex is surrounded by DotG, which is another essential Dot/Icm protein spanning from the outer membrane to the inner membrane (Figs. 1D and 1E).

### The outer membrane core complex (OMCC) is associated with three distinct porin arrays

The *in-situ* OMCC structure is largely consistent with a previous cryo-EM structure of the purified OMCC(32, 33) but with notable conformational differences. Compared to the purified OMCC, the inner diameters of the OMC and PR *in situ* are about 6-8 Å smaller, resulting in a more compact OMCC structure (SI Appendix, Fig. S5). Notably, the *in-situ* dome structure – formed by the C-terminal residues of DotG (residues 862–1046) – is resolved at 3.08 Å resolution. The dome appears to be more stable in the native membrane than in the purified OMCC(32, 33).

We also identified an uncharacterized density between Dis3 proteins in the OMCC (SI Appendix, Fig. S6A). Utilizing Modelangelo(37), a machine-learning–based atomic modeling tool, combined with Dali structural homology searches, we determined that this density corresponds to a small α/β-structured protein (73 residues) encoded by an uncharacterized gene *(uniport:A0AAN5PM39)* (SI Appendix, Fig. S6B-S6E). We named this protein Dis4, in line with the previously described species-specific subunits Dis1, Dis2, and Dis3. Dis4 interacts with the C-terminus of DotC in an interdigitated manner, whereby the C-terminus of DotC inserts into the middle of Dis4, forming a 3-stranded β-sheet structure (SI Appendix, Fig. S6C). Given its intimate association with DotC and other components of the OMCC, Dis4 is likely critical for initial assembly of the Dot/Icm machine while the exact role of Dis4 remains to be determined.

Although Dot/Icm machines have previously been shown to co-purify with MOMPs(38, 39), it is quite striking to observe three distinct porin proteins – here named Porins I, II, and III – closely associated with the OMCC in our *in-situ* structure (Figs. 1C-1E and 2A-2C). Porins I and II each form trimers and assemble as two concentric rings with C13 symmetry around the periphery of the OMCC (Figs. 2B and 2D). Based on structural comparison with all β-porin-like proteins encoded in the *L. pneumophila* genome, Porin I is a MOMP(40) likely encoded by *lpg2961* (Fig. 2D; SI Appendix, Fig. S7A). It adopts a 10-stranded β-barrel structure and interacts specifically with the α-helix 1 and α-helix 2 of Dis1 in the OMCC (Figs. 2E and 2F). Porin II is located above Dis3 at the distal periphery of the OMCC. The overall features of Porin II are consistent with those of Porin I, suggesting it may be encoded by *lpg2961* (SI Appendix, Fig. S3F and S3G). To verify the role of Lpg2961 in forming the Porin I and II arrays, we constructed a lpg2961 transposon insertion mutant (lpg2961::Tn) to disrupt protein expression and determined the *in-situ* structures of the Dot/Icm machine by cryo-ET and subtomogram averaging. In the resulting structure, the densities corresponding to Porin I and II were completely absent or partially absent (Figs. 2G-2J), implying that lpg2961 and other similar porin genes are likely involved in formation of the Porin I and II arrays. Different from Porins I and II, Porin III forms a ring with C16 symmetry tightly encircling the dome near the exit gate of the Dot/Icm machine (Fig. 2C). Porin III is a small MOMP with an 8- stranded β-barrel and a broad extracellular domain although the precise sequence remains unknown. Notably, these three porin arrays protrude from the lipid bilayer and extend approximately 25 Å into the extracellular space (Fig. 2), suggesting plausible roles in host-pathogen interactions during infection.

**Figure 2.**
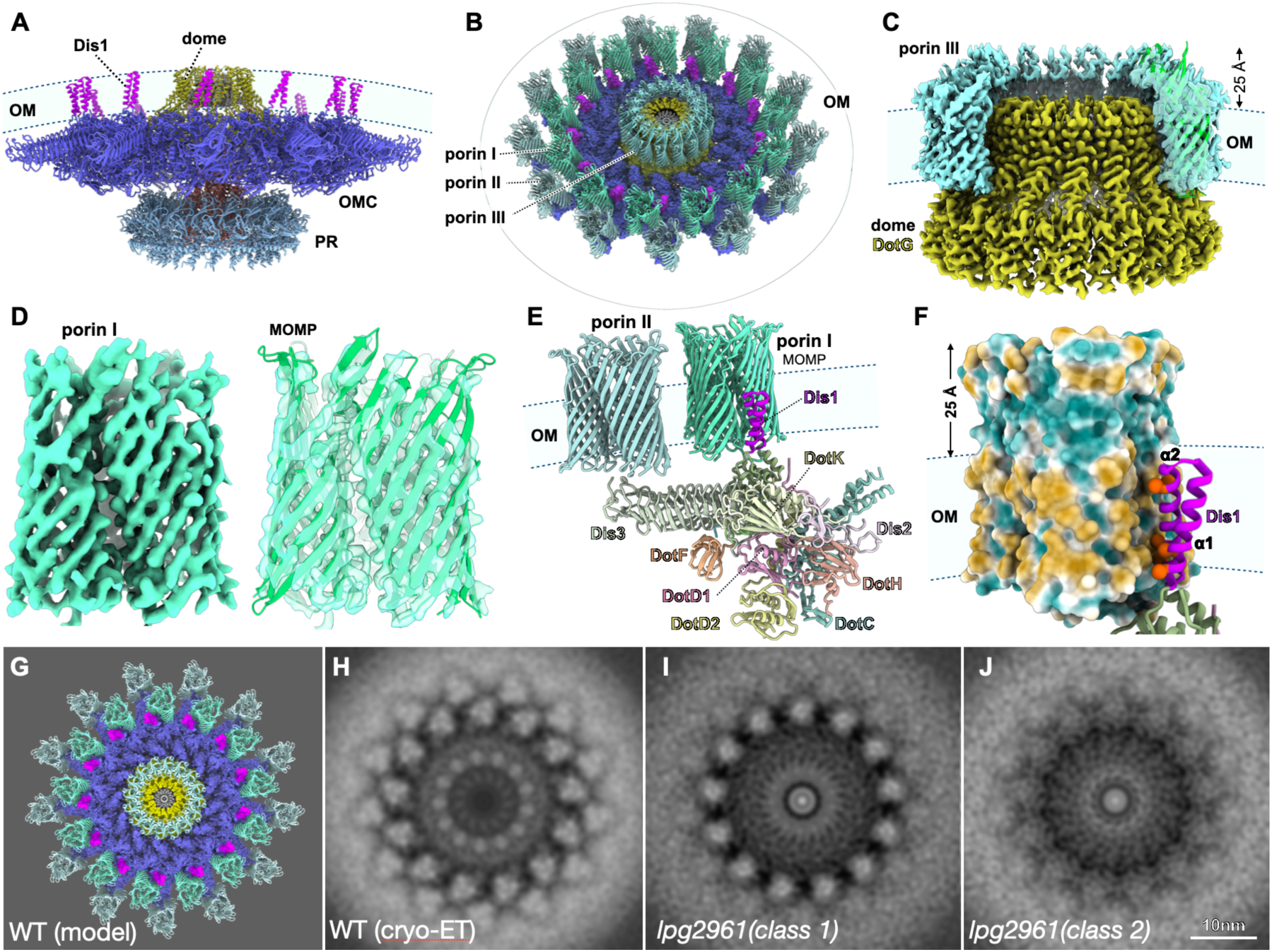
The outer membrane core complex (OMCC) is associated with three porin arrays in the outer membrane. (**A**) The structure of OMCC consisting of the disk-shaped outer membrane cap (OMC), dome, and periplasmic ring (PR). The dome and transmembrane helices (α1 and α2, colored by purple) of Dis1 are embedded in the outer membrane (OM). (**B**) A tilt view shows the integration of three porin arrays with the OMCC. Porin I is adjacent to the transmembrane helices of Dis1, Porin II is above the distal arm of OMC, and Porin III forms a ring closely wrapping around the dome (yellow). (**C**) A close-up view of the Porin III ring around the dome formed by residues 842–1046 of DotG. The Porin III extends approximately 25 Å into the extracellular space. (**D**) Refined cryo-EM map of the Porin I trimer and model of a MOMP likely encoded by *lpg2961*. (**E**) A close-up view of Porin I and Porin II integrated with the OMCC. The subunits of the OMCC are colored differently. Porin I interacts specifically with the transmembrane helices of Dis1 (purple), and Porin II is adjacent to the distal region of Dis3 (yellow). (**F**) Surface view of the Porin I trimer reveals the hydrophobic surface embedded in the OM and the interactions with two transmembrane helices of Dis1. (**G**) A top view of the OMCC and its association with porin arrays. (H) A section of the wild-type Dot/Icm machine reconstruction shows the Porin I and II trimers with C13 symmetry by cryo-ET. Two class averages from the *Ipg2961*::Tn mutant show that the Dot/Icm machines lacking Porin II array (I) and lacking Porin I/II arrays (J).

### DotA and IcmX form a protochannel in a closed conformation

IcmX forms a pentamer in the central axis of the OMCC. Each IcmX monomer, consisting of 466 amino acids, is structurally divided into N- and C-terminal α-helices and extended domains (residues 90–290) (Fig. 3A; SI Appendix, Fig. S8A and S9). The five N-terminal helices of the IcmX pentamer assemble into a bundle resembling a pore-forming structure (Fig. 3B; SI Appendix, Fig. S9). The extended domains (residues 90–290) create a loop-rich surface that significantly enlarges the overall size of the IcmX pentamer, likely facilitating its interactions with the OMCC (Fig. 3B).

**Figure 3.**
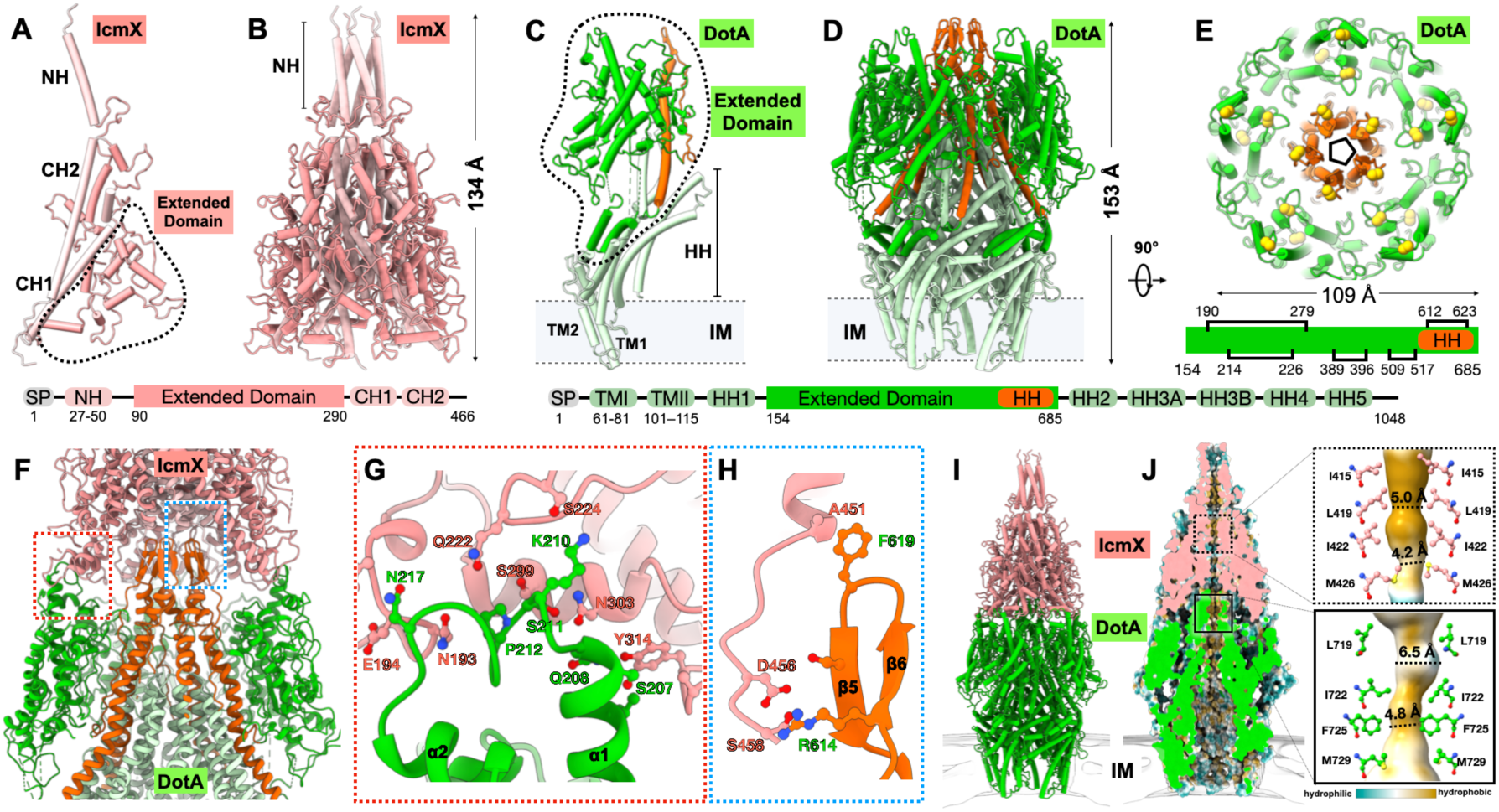
*In-situ* structures of DotA and IcmX. (**A, B**) IcmX contains a SP and structurally conserved α-helices at its N- and C-termini (NH and CH) and an extended domain. Monomeric and pentameric structures of IcmX are shown as cylinder diagrams. The conserved N- and C-terminal helices (NH and CH) are colored light red, and the extended domain is highlighted in red. (**C**, **D**) DotA contains a signal peptide (SP), Two conserved N-terminal transmembrane helices (TMI and TMII), five hydrophobic helices (HH1-HH5), and a large extended domain inserted between them. Monomeric and pentameric structures of DotA are shown as cylinder diagrams. TM1, TM2, and hydrophobic helices (HH) are colored light green. The extended domain is colored in green, with one hydrophobic helix and a β-hairpin highlighted in orange. (**E**) A top view of the extended domains of DotA. Each unit contains five disulfide bonds, depicted as yellow spheres with the involved cystine residues shown below. (**F**) Extensive interactions between IcmX and DotA are primarily mediated by their extended domains. The detailed interactions are shown in two panels (**G, H**). (**I**) An overall structure of the DotA-IcmX complex. (**J**) A cross-sectional view of the lumen of the DotA- IcmX protochannel. Two enlarged views of the IcmX and DotA residues in the lumens.

Underneath the IcmX complex, DotA forms a pentamer anchored in the inner membrane (Figs. 3C, 3D; SI Appendix, Fig. S8B, S10, S11). Each DotA protomer features two N-terminal helices (TM1 and TM2) embedded in the inner membrane and five long hydrophobic helices (HH1-HH5) (Fig. 3C; SI Appendix, Fig. S10). Pentamerization of DotA creates a large, hollow chamber in the inner membrane and periplasm (Fig. 3D; SI Appendix, Fig. S11A-S11D). DotA contains a large hydrophilic extended domain (residues 154–685) between the hydrophobic helices (Figs. 3C, 3D; SI Appendix, Fig. S10 and S11). This extended domain of DotA adopts a unique fold with no homologous structure identified through Dali or Foldseek searches(41, 42), but it is conserved among DotA family proteins in *Legionella* species(43). This unique domain is characterized by a core of crossing helices (α1, α4, and α5) flanked by small β-sheets and rich loop regions (SI Appendix, Fig. S10). The extended domain connects to the chamber through a unique parallel helix-loop (helix α11 and the loop between α10 and β5), anchoring it like tiles on the surface of the chamber (SI Appendix, Fig. S10). Two short helices (α6 and α7) insert into the midsection of the chamber, further stabilizing the structural assembly. Moreover, the extended domain of DotA is enriched with five potential disulfide bonds per protomer surrounding the top of the chamber (Fig. 3E; SI Appendix, Fig. S10D).

DotA and IcmX interact primarily through their extended domains (Fig. 3F). Within DotA, a unique pentameric β-hairpin inserts into the base of the IcmX pentamer. This insertion is complemented by an extensive peripheral interface, forming a shape-matched connection that interlocks the two proteins (Fig. 3F). Their interaction surface area exceeds 4,000 Å² and is mediated by hydrophilic interactions involving side chains and main chains between the loop (residues 205–220) of DotA and base of the IcmX extended domain (Figs. 3G and 3H). This interaction pattern is strikingly distinct from the VirB5-VirB6 complex in the R388 T4SS (SI Appendix, Fig. S12), where the interface primarily serves as a platform for pilus assembly(44, 45). Given the absence of pilus-related genes from the Dot/Icm operon in *L. pneumophila* and the lack of observed pilus structures in the Dot/Icm system, this structural divergence may reflect an alternative mechanism for effector protein translocation in the absence of a pilus conduit.

Notably, the DotA-IcmX complex exhibits a hydrophobic interior along its central axis (Figs. 3I and 3J). At the top of the DotA chamber, hydrophobic residues (L719, I722, F725, and M729) form a narrow, neck-like structure with an inner diameter ranging from ∼4.8 to 6.5 Å (Fig. 4J). Similarly, the central lumen of the IcmX complex is also constricted, measuring 4.2 to 5.0 Å (Fig. 4J). In contrast to the previous observation suggesting that IcmX forms a plug(46), our findings indicate that IcmX and DotA assemble into a pentameric protochannel with a lumen narrow enough to prevent substrate translocation, thereby maintaining the Dot/Icm machine in an inactive state. This closed conformation is further stabilized by extensive interactions between the extended domains of DotA and IcmX. This architecture of the Dot/Icm machine is consistent with the notion that the system remains in an inactive state preventing the premature secretion of effectors until close contact is established with a recipient host(23).

**Figure 4.**
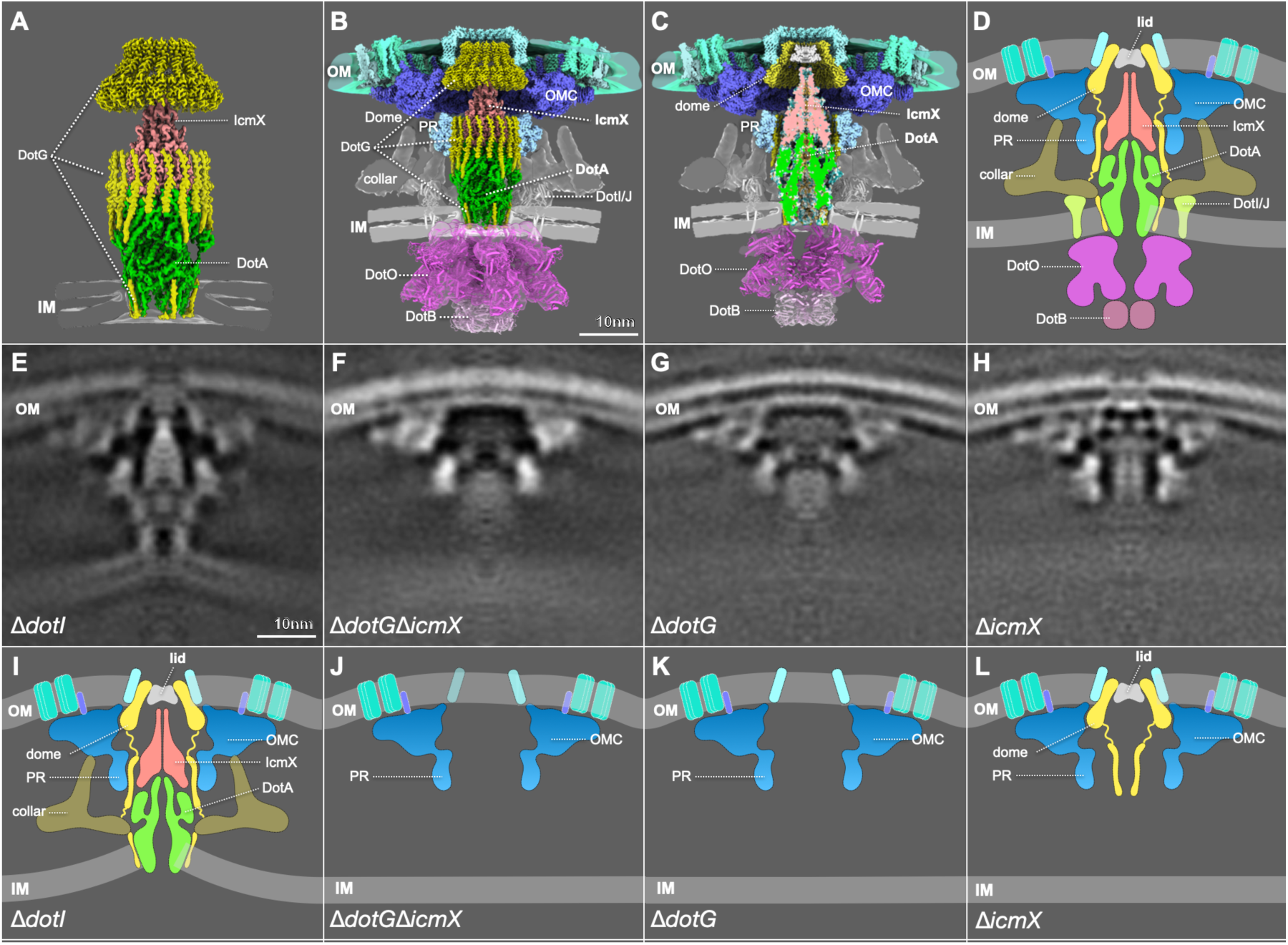
Assembly of the DotA-IcmX protochannel in the intact Dot/Icm machine. (**A**) The *in-situ* structure reveals that DotG assembles into a cage-like structure that encases the DotA-IcmX protochannel. DotG is depicted in yellow, while DotA and IcmX are colored green and red, respectively. (**B**) The overall structure of the intact Dot/Icm machine including atomic models of the OMCC and DotA/IcmX complex, the cytoplasmic ATPase complexes (DotO and DotB), the arche- like structure formed by DotI and DotJ and the collar density. (**C**) A cross-sectional view of the DotA/IcmX protochannel showing the central lumen, indicating that the Dot/Icm machine is maintained in an inactive state, preventing substrate translocation. (**D**) A schematic model of the Dot/Icm machine in its inactive state. (**E-H**) Cross-sectional views of the cryo-ET structure of Dot/Icm machine from Δ*dotI*, Δ*dotG*Δ*icmX*, Δ*dotG*, and Δ*icmX*, respectively. (**I**) A schematic model of the Dot/Icm machine lacking DotO/DotB ATPase complexes in the Δ*dotI* mutant. (**J**) A schematic model of the Dot/Icm machine lacking DotG, DotA/IcmX, DotO/DotB ATPase complexes in the Δ*dotG*Δ*icmX* mutant. (**K**) A schematic model of the Dot/Icm machine lacking DotG, DotA/IcmX, DotO/DotB ATPase complexes in the Δ*dotG* mutant. (**L**) A schematic model of the Dot/Icm machine lacking DotA-IcmX, DotO-DotB ATPase complexes in the Δ*IcmX* mutant. The dome of DotG remains while the rest of DotG is not well resolved.

### DotG is critical for assembly of the DotA-IcmX protochannel

Our *in-situ* structure resolves three segments of DotG, illustrating its organization around the DotA- IcmX complex and spanning both the inner and outer membranes (Fig. 4A). The first segment (residues 2–42) comprises the N-terminal transmembrane helices, positioned adjacent to the transmembrane helices TMI and TMII of DotA and exhibiting C5 symmetry (Fig. 4A; SI Appendix, Fig. S13). The second segment (residues 791–824) forms helices that constitute the interior wall of the PR. The third segment (residues 862–1046) forms a dome structure with C16 symmetry within the outer membrane (Fig. 4A). While the majority of DotG (total 1048 residues), particularly the pentapeptide repeat motif in the middle domain, remains unresolved in our cryo-EM structure, the resolved regions suggest that DotG assembles into a cage-like structure encasing the DotA-IcmX protochannel (Fig. 4A).

To better understand the intact structure of the Dot/Icm machine and assembly of the DotA-IcmX protochannel within, we employed cryo-ET and subtomogram averaging to analyze approximately 2,923 Dot/Icm particles in wild-type bacteria. This approach enabled us to resolve the flexible collar-like densities and small arch-like densities around the DotA-IcmX complex in the periplasm as well as the cytoplasmic ATPase structures at 16Å resolution (Figs. 4B-4D; SI Appendix, Fig. S14). Leveraging AlphaFold prediction and our cryo-EM/ET maps, we built a more complete model of the Dot/Icm machine, including two ATPase complexes (DotB and DotO) and inner membrane proteins (DotI and DotJ) (SI Appendix, Fig. S14A-S14I; SI Appendix, Movie S1). Consistent with our previous reports(31, 34), the DotO complex forms a large cytoplasmic channel, functioning as a conduit for substrates to access the chamber formed by DotA as well as providing the necessary translocation energy. However, the DotA-IcmX protochannel prevents the substrate translocation due to the narrow hydrophobic lumen along its central axis (Figs. 4B-4D).

To validate the model, we utilized cryo-ET and subtomogram averaging to analyze mutants lacking DotI, DotG, IcmX, and IcmX/DotG, respectively. Our cryo-ET data revealed that DotI is critical for assembly of the cytoplasmic ATPase complexes but is not required for assembly of the DotA-IcmX protochannel or DotG cage/collar (Figs. 4E and 4I). By contrast, DotG and IcmX are indispensable to each other as both the DotG cage/collar and DotA-IcmX protochannel fail to assemble in the absence of either DotG or IcmX (Figs. 4F-4L).

### Dot/Icm machines undergo major conformational changes upon host contact

T4SSs translocate substrates in response to activating signals conveyed upon contact with recipient cells(22, 47, 48). The dynamic nature of effector translocation has historically made it challenging to capture active conformations of T4SSs, including those of the Dot/Icm machines in *L. pneumophila*. Consequently, mechanistic details of how T4SSs deliver effector proteins into host cells have remained poorly understood. To capture active states in the Dot/Icm machine, we used cryo-ET to visualize cryo*-FIB-milled lamellae* of macrophages infected with *L. pneumophila*. These images reveal some Dot/Icm machines attached to the membrane of the LCV (Figs. 5B-5D; SI Appendix, Movie S2) by a short tether, supporting an activation mechanism by which host membrane contact enables formation of the tether and direct translocation of effector proteins into recipient cells(48). However, due to the challenge of the cryo-FIB-ET approach, structural details at the interface between the Dot/Icm machine and host membrane remained limited.

**Figure 5.**
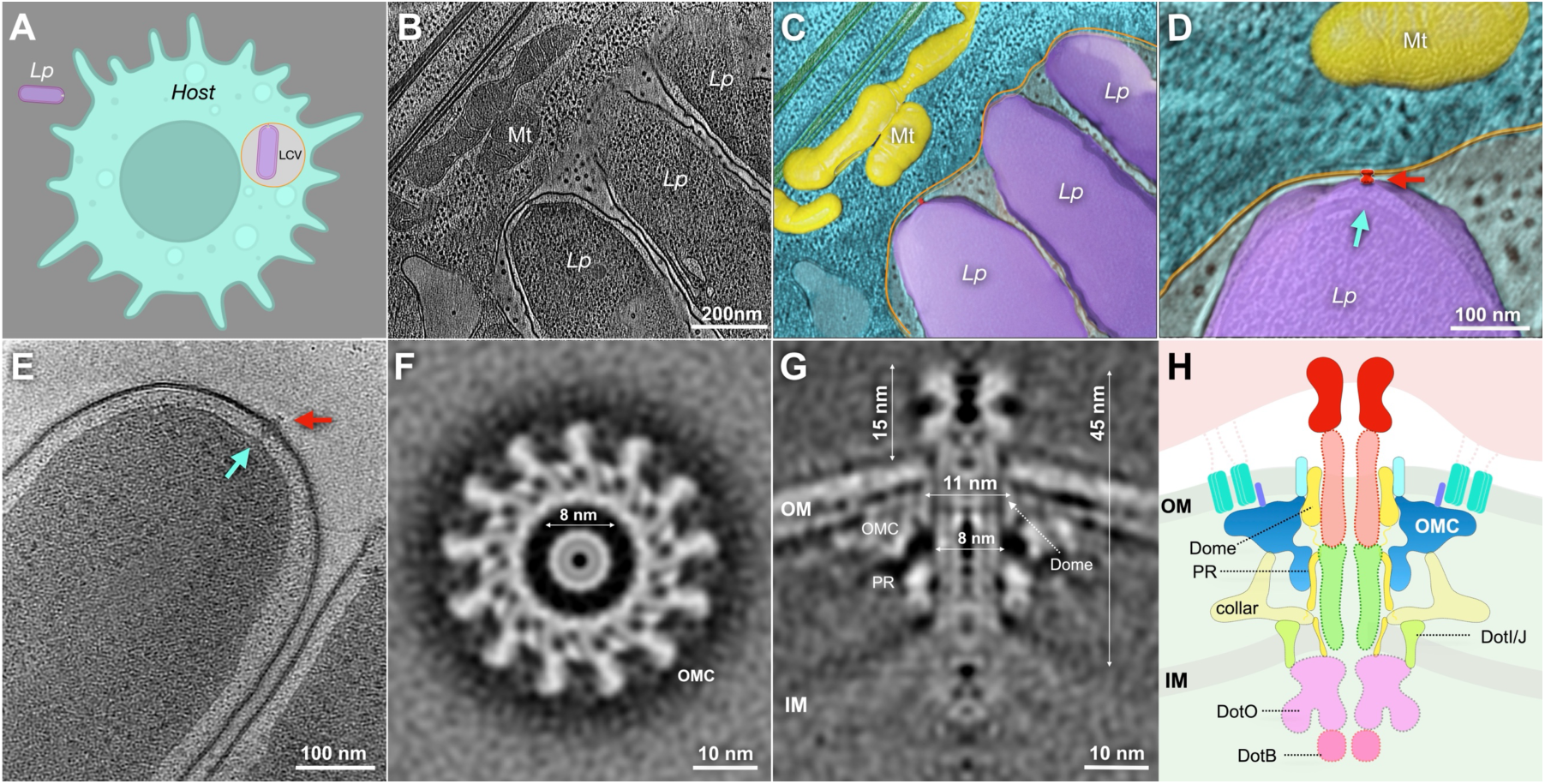
The DotA-Icm complex undergoes extensive conformational changes upon host contact. **(A)** A schematic model of LCV in a host cell. (**B**) A cryo-ET image of *L. pneumophila* (*Lp*) cells inside the LCV. Mt, mitochondrion. (**C, D**) 3D segmentation of multiple *L. pneumophila* cells attached to the membrane of the LCV through a short tether (red arrow). (**E**) A cryo-ET snapshot of a cell pole after incubation with macrophage lysate shows a Dot/Icm machine (blue) with an extension (red). (**F**, **G**) Top and cross-sectional view of the cryo-ET structure of the Dot/Icm machine after incubation with lysate. Notably, a novel transenvelope channel with a length of 45 nm and a diameter of 8 nm and a distinct 15 nm extracellular protrusion emerges upon activation. (**H**) A schematic model of the Dot/Icm machine in an active state.

To overcome this challenge, we developed an alternative approach to increase activation frequency of the Dot/Icm machines by incubating *L. pneumophila* cells in a macrophage lysate containing membrane vesicles for 30 minutes. As the lysate contains various host factors that could activate the Dot/Icm machines to translocate effector proteins, this approach allowed us to capture a distinct structural phenotype in which an additional density extends from the Dot/Icm machine into the extracellular space (Fig. 5E; SI Appendix, Movie S3). Subsequently, subtomogram averaging of these particles revealed considerable conformational changes compared to the inactive state of the Dot/Icm machine. The DotA-IcmX complex likely transitions from a closed protochannel to an extended transenvelope channel with a widened lumen. By contrast, the OMC remains structurally similar to what is observed in the inactive state, while the dome region undergoes partial opening from 8 nm to 11 nm, accommodating formation of the transenvelope channel 45 nm in length and 8 nm in diameter (Figs. 5G and 5H). Notably, a distinct 15 nm extracellular protrusion emerges upon activation. The length of the protrusion aligns with the tether observed between *L. pneumophila* and the LCV membrane, suggesting that the structure facilitates membrane engagement and effector delivery from the bacterial cytoplasm into host cells (Fig. 5H). These findings provide critical insights into the dynamic structural rearrangements underlying T4SS activation, advancing our understanding of these versatile secretion systems.

## Discussion

The Dot/Icm T4SS of *L. pneumophila* is one of the most complex and versatile machines in the bacterial kingdom. Over 27 different proteins are involved in assembly and activation of the Dot/Icm T4SS. Though the system has been extensively characterized since its discovery in 1998, the molecular mechanisms underlying its assembly and activation have remained incomplete, largely due to limited structural information of the intact machine in its functional state. To overcome this limitation, we developed a highly innovative approach combining *in-situ* cryo-EM and cryo-ET with molecular genetics. Notably, *in-situ* cryo-EM enabled us to reveal at near-atomic resolution three novel porin arrays intimately associated with the OMCC and a DotA-IcmX pentamer at the central axis of the Dot/Icm machine. Cryo-ET enabled us to uncover the overall architecture of the intact Dot/Icm machine and its major conformational changes upon activation, laying a foundation for mechanistic studies of Dot/Icm assembly and activation.

A central finding of our studies is that DotA and IcmX form a protochannel within the Dot/Icm machine. Recent reports based on deletion phenotypes or the homology of type VI secretion system (T6SS) components had suggested that DotG or IcmF forms the periplasmic channel-like density and that IcmX functions as a plug(46, 49). However, our *in-situ* structures clearly demonstrate that DotA and IcmX form a protochannel with a narrow lumen in closed conformation. Remarkably, the DotA-IcmX protochannel is surrounded by DotG, which forms a cage-like structure spanning the outer and inner membrane and may play a role in regulating the DotA-IcmX protochannel. Therefore, our results solve a longstanding question as well as provide new insights into why these three proteins are required for assembly and function of the Dot/Icm machine.

Crucially, we capture the active state of the Dot/Icm machine, in which we propose that the protochannel of DotA-IcmX undergoes a dramatic conformational change to form a ∼45 nm transenvelope channel facilitating host membrane attachment and translocation of effector proteins from the cytoplasmic space into recipient cells. Our data suggest that, upon system activation, the conformational change of the DotA pentamer pushes the IcmX pentamer outward, forming a conduit that spans the bacterial envelope and ultimately engages the host membrane. This conduit serves as the essential interface for effector protein translocation, directly connecting the bacterial cytoplasm to the host cell environment. Our discovery of the extracellular conduit provides new insights into the mechanism of T4SS-mediated host-pathogen interactions, suggesting a previously unrecognized structural adaptation facilitating direct molecular communication during infection.

We thus propose a “sewing machine” model for type IV secretion, whereby the central DotA-IcmX channel functions as a dynamic machine stabilized by the OMCC and periplasmic cage (SI Appendix, Movie S4). By expanding, the DotA-IcmX complex would deliver protein threads into a host cell (SI Appendix, Fig. S15; SI Appendix, Movie S4). One consequence of the dynamic nature of DotA-IcmX assembly may be that this complex is occasionally ejected by the Dot/Icm machine, which would explain why DotA and IcmX remain the only two Dot/Icm proteins known to be secreted into *L. pneumophila* culture supernatants by a process requiring a functional Dot/Icm apparatus(50, 51). Our findings advance understanding of the complex architecture and activation mechanisms of the Dot/Icm T4SS, providing a molecular framework for exploring its physiological roles and therapeutic potential. Our studies also lay a foundation for investigations aimed at elucidating the molecular composition, assembly process, and functional mechanisms of T4SSs.

## Materials and Methods

### Bacterial strains

The bacterial strains used in this study were derived from *Legionella pneumophila* LP01 and LP02. A DotB E191K mutant strain (without GFP tags), previously characterized for its ability to stably bind DotO without ATP hydrolysis activity and exhibited a more population intact assembly of the Dot/Icm machine(31), was utilized for *in-situ* cryo-EM single-particle analysis. For cryo-ET analysis, we generated a series of mutants, including single and double deletions of key Dot/Icm components: Δ*dotI*, Δ*icmX* and Δ*icmXdotG*. Mutants were constructed using standard recombinant DNA techniques as described previously(31). Additionally, to investigate the candidate porin proteins (*lpg2961*), we generated transposon insertion mutants in *lpg2961* at the residue A167 between F168 (lpg2961::Tn) using established protocols(52). The mutants were validated by PCR and DNA sequencing to confirm accurate insertion sites. *L. pneumophila* cells were cultured on charcoal yeast extract (CYE) agar plates at 37°C.

### Preparation of frozen-hydrated specimens

For cryo-EM grid preparation, *L. pneumophila* cells were revived form -80° on CYE plates at 37°C for 5 days, re-streaked onto fresh plates, and incubated for an additional 48 hours. Bacterial suspensions in PBS (OD600 ≈1.5) were applied to glow-discharged Quantifoil Cu 2/1 300 mesh grids for 1 minute. Grids were blotted for 6–8 seconds under 90% humidity and rapidly plunge-frozen in liquid ethane using a GP2 plunger (Leica).

### Cryo-EM data collection and preprocessing

Data acquisition was performed on a Titan Krios electron microscope (Thermo Fisher Scientific) operated at 300 kV, equipped with a K3 direct electron detector and a Gif energy filter (Gatan) in super-resolution mode (pixel size: 0.534 Å). SerialEM(53) was used to facilitate multi-shot acquisition targeting cell poles within ice holes. Images were recorded with a total dose of 73 e⁻/Å² distributed over 40 frames (0.07 s/frame), with defocus values ranging from −1.0 to −2.0 μm (SI Appendix, Table S1). Motion correction and contrast transfer function (CTF) estimation were performed in CryoSPARC(54, 55).

### *In-situ* single particle selection using GisSPA and Topaz

Traditional particle-selection methods proved inadequate due to high cellular background noise and sparse localization of Dot/Icm complexes at bacterial poles. To address this, rotational-translational template matching was implemented using GisSPA (GPU-accelerated in situ single-particle analysis)(35), detection for *in-situ* particle localization (SI Appendix, Fig. S1). A low-pass filtered (8 Å) and detergent-masked template of the outer membrane core complex (EMD-24004) was applied to 1024×1024-pixel cropped small micrographs (2×binned). Detection thresholds were set at a correlation coefficient of 6.5, with Euler angles sampled at 3° intervals. Local peaks within 30 pixels (unbinned) were clustered, yielding 2,080 initial particles from 2,857 micrographs. Coordinates and orientation parameters were exported to RELION v3.1 for particle extraction(56). Subsequent 2D classification retained particles displaying Channel (DotA) densities and membrane features, followed by 3D classification without alignment to refine the particle selection.

Topaz(36) was iteratively trained using GisSPA-derived coordinates, with a particle diameter parameter of 350 Å. This approach enabled identification of ∼10 particles per micrograph, enhancing coverage of the Dot/Icm complexes in the cell poles.

### Image processing and structure determination

All structural analyses were performed using CryoSPARC(54, 55), with the overall workflow summarized in SI Appendix, Fig. S2. Initial particle selection identified 222,686 potential particles, which were extracted from a total of 49,978 micrographs and refined using a 612-pixel box size (1.335 Å/pixel). Homogeneous refinement with C13 symmetry was first applied to the dataset using an initial model generated by GisSPA(35), yielding an overall reconstruction at 6.64 Å resolution. Subsequent iterative heterogeneous refinement and 3D classification were conducted to improve particle selection, resulting in a high-quality subset of 76,409 particles.

For the refinement of the OMCC, local refinement combined with local CTF correction was applied to the selected subset using C13 symmetry, achieving a final resolution of 2.96 Å. To further resolve the PR, signal subtraction was performed on OMCC-containing particles using a soft mask, followed by C13 symmetry expansion. The expanded particles were subjected to 3D classification without alignment to identify structural features consistent with C18 symmetry. After selecting classes with clear PR structural features and removing duplicate particles, local refinement with C18 symmetry was performed, resulting in a 3.04 Å reconstruction.

This strategy was extended to the DotA-IcmX and dome sub-complexes using C5 and C16 symmetries, respectively (SI Appendix, Fig. S2). Since the channel component does not assemble stably in all Dot/Icm machines, only 37,503 particles (49% of the OMCC particles) exhibited high-quality channel density, leading to a 3.63 Å reconstruction with C5 symmetry.

To analyze the interaction between the PR and the channel, a symmetry breaking reconstruction of the PR-channel complex was performed. C5-expanded particles from the 3.63 Å channel reconstruction underwent 3D classification without imposed symmetry. Particles exhibiting both PR and channel features were selected for subsequent local refinement with C1 symmetry, yielding a 4.63 Å reconstruction.

For porin protein structure determination, Porins I and II were reconstructed using C13- expanded particles by shifting the refinement center to porin positions, followed by local C1 refinement. Porin III was resolved through dome-focused reconstruction using C16 symmetry.

### Model building and refinement

For the OMCC, the atomic model (PDB-7MUD) was initially docked into the cryo-EM density maps using ChimeraX(57). Individual components were manually adjusted in Coot v0.8.9.1(58) to optimize their fit within the map. Model refinement was performed iteratively using Phenix 1.21(59) to improve residue alignment with the density map.

The IcmX model was predicted using AlphaFold3(60) as an initial template. Subsequent manual adjustments were made in Coot to enhance its fit within the cryo-EM map. Since the identity of the channel-forming protein remained unknown, we employed ModelAngelo(61) to generate a model using *L. pneumophila* (ATCC 33152) genome-derived protein sequences. Although the sequence assignment was not fully accurate, the predicted secondary structure and overall fold were well-matched to the cryo-EM density.

To further validate the channel-forming protein identity, we performed structural homology searches using a locally installed DALI server(62). This analysis compared our density-based model against a construct library of *L. pneumophila* AlphaFold-predicted proteins. The highest-scoring match was identified as DotA, which was then manually refined in Coot to ensure proper residue placement within the density map.

### Porin sequence identification and structural modeling

To identify potential porin candidates, we screened all β-barrel-like proteins from the *L. pneumophila* AlphaFold-predicted structural library. A total of 25 candidate proteins were identified, and monomeric and trimeric models were generated for structural fitting and comparison.

For Porin I, five candidate proteins exhibited 10 β-stranded structural features, with three showing approximate structural compatibility based on overall size and β-sheet organization. Among them, *Lpg2961* (UniProt: Q5ZRCI) provided the best fit. The density map of Porin II exhibited a size and shape similar to Porin I. Thus, we hypothesize that Porin II may represent a structurally similar or identical protein.

Final model refinement was conducted through iterative adjustments in Phenix, ensuring optimal residue alignment within the cryo-EM density map. Model geometry was validated using MolProbity(63) (SI Appendix, Table S2). Local resolution was estimated with FSC 0.143 criterion in CryoSPARC(54, 55).

### *L. pneumophila*-infected J774 macrophages for cryo-FIB milling

J774 macrophages were cultured on Quantifoil R2/1 gold EM grids placed in 35-mm glass-bottom dishes. Grids were sterilized by UV-exposed ethanol immersion (30 min) followed by four PBS washes, then preconditioned overnight in DMEM (Gibco) supplemented with 10% heat-inactivated FBS (Gibco). Cells were seeded at 5 × 10⁶ cells/dish and incubated for 48 hr. After PBS washing, cells were serum-starved in DMEM for 2 hr prior to infection. Wild-type *L. pneumophila* grown on CYE agar were PBS-washed and used to infect macrophages at MOI 50 for 10 hr. Grids were plunge-frozen using a manual plunger, transferred under liquid nitrogen to a cryo-FIB system, and sputter-coated with a 1 kV/15 mA platinum layer (15 s) followed by organometallic platinum deposition (GIS). Lamellae were milled using a 30 kV gallium ion beam at 14° tilt, progressively reducing beam current to achieve ≤200 nm thickness. 12 tilt series were acquired at ×15,000 magnification with a Volta phase plate (VPP) for contrast enhancement. 22 Dot/Icm machines were detected in the FIB data. Three of them appear tethering to the LCV membrane.

### Macrophage lysate-induced Dot/Icm activation for cryo-ET grid preparation

THP-1 human monocytic cells were differentiated into macrophage-like cells by incubation with 100 nM phorbol 12-myristate 13-acetate (PMA, Sigma-Aldrich) in RPMI 1640 medium (Gibco) supplemented with 10% fetal bovine serum (FBS, Gibco) at a density of 1 × 10⁶ cells/mL. Cells were incubated for 48 hours at 37°C in a humidified atmosphere containing 5% CO₂ to promote differentiation. Following differentiation, non-adherent cells were removed by washing with phosphate-buffered saline (PBS, pH 7.4), and the adherent cells were replenished with fresh RPMI medium containing 10% FBS. Cells were then incubated for an additional 48 hours to achieve full differentiation. Differentiated macrophage-like cells were detached using a cell scraper, collected, and resuspended in PBS at a final concentration of 2 × 10⁷ cells/mL. Cell lysis was performed by mechanical disruption using 2 mm glass beads (Sigma-Aldrich), with continuous vortexing for 10 minutes at room temperature. The resulting lysates were clarified by centrifugation at 1,000 × g for 10 minutes at 4°C to remove cell debris. The supernatant was subsequently filtered through 0.45 μm syringe filter to obtain clarified cell-free lysate. Wild-type *L. pneumophila* (OD600 1.0 in PBS) were incubated with the prepared THP-1 cell lysate at a 1:1 volume ratio. Incubations were performed at 37°C for three different time points: 30 minutes, 1 hour, and 2 hours, with gentle agitation to facilitate interactions. For cryo-ET, each sample from the three incubation time points was mixed with 10 nm gold fiducials and applied to glow-discharged Quantifoil R2/1 Cu 200 mesh grids and vitrified using a GP2 plunge freezer (Leica).

### Cryo-ET data acquisition and subtomogram averaging

Tilt series were collected on a Titan Krios electron microscope (Thermo Fisher Scientific) operated at 300 kV, equipped with a K3 direct electron detector and a Gif energy filter (Gatan) in super-resolution mode (pixel size: 1.06 Å) using SerialEM(53). Tilt ranges span from -48° to +48° with 3° increments (total dose ∼70 e⁻/Å², defocus -4.8 μm). Recorded images were initially motion-corrected using MotionCorr2(64) and subsequently stacked by IMOD(65). Tilt series alignment was performed using fiducial markers or fiducial-free alignment by IMOD. After CTF estimation (Gctf)(66) and phase flipping (IMOD ctfphaseflip), Tomo3D(67) was used to generate tomograms either simultaneous iterative reconstruction technique (SIRT) or weighted back-projection (WBP). SIRT reconstructions were utilized for particle selection and WBP tomograms were used for subtomogram averaging. The number of tomograms and particles utilized for each strain is detailed in SI Appendix, Table S3.

The i3 package was used to perform 3D alignment and classification as described previously(68). Initially, Dot/Icm machine were aligned within 8× binned tomograms based on the initial orientation. Subsequently, subtomograms were extracted from unbinned tomograms using the refined positions and 4× binned for alignment. This process was reiterated with 2× binned subtomograms to enhance structure detail. Fourier shell correlation calculated by comparing two randomly divided halves of the aligned images was used to estimate *in-situ* structure resolution.

### Building model into cryo-ET maps

Due to the resolution limitations of our cryo-ET map, we employed AlphaFold3 structural predictions for DotO, DotI, and DotJ to assistant in model building. Given the homology of these components to high resolution structure (PDB:8RTB) of VirB4 and VirB8 in the R388 T4SS and supporting evidence of their hexameric assembly in the inner membrane, we constructed a predicted model based on a DotO:DotI:DotJ stoichiometry of 2:2:1. The model was then fitted into the cryo-ET density map and manually adjusted using Coot to optimize alignment with the density. C6 symmetry was subsequently applied to generate the hexameric structures of the DotO-DotB complex and DotI-DotJ complex.

## Acknowledgments

This work was funded by National Institutes of Health (NIH) grant R01AI152421 to Jun Liu and Craig R. Roy. We thank Jing Cheng for advice in using GisSPA. We thank Jennifer Aronson for critical reading of the manuscript. Cryo-EM data were collected at the Yale Cryo-EM resources funded in part by the NIH grant 1S10OD023603-01A1.

## Data Availability

All cryo-EM maps and atomic coordinates for this study have been deposited in the Electron Microscopy Data Bank (EMDB) and the Protein Data Bank (PDB). These include locally refined maps of the disk-shaped outer membrane cap (OMC) under accession codes EMD-49393 (PDB 9NGU), the periplasmic ring (PR) under EMD-49394 (PDB 9NGV), the dome under EMD-49395 (PDB 9NGW), the DotA-IcmX complex under EMD-49396 (PDB 9NGY), the PR-DotA-IcmX complex at C1 symmetry under EMD-49398 (PDB 9NH0), Porin I under EMD-49399 (PDB 9NH1), and Porin III under EMD-49395 (PDB 9NH2).

## Author Contributions

Conceptualization, J.Y., C.R.R., and J.L.; methodology, J.Y., S.H., D.P., D.C., S.T., W.G., J.M.B., S. W., and J. L.; investigation, J.Y., S.H., D.P., D.C., S.T., J.M.B., C.R.R., and J. L.; visualization, J.Y., S.H., J.M.B.; formal analysis, J.Y., S.H., D.P., D.C., S.T., and J.L.; funding acquisition, C.R.R., and J.L.; supervision, C.R.R., and J.L.; writing – original draft, J.Y., S. H., and J.L.; writing – review and editing, J.Y., S.H., D.P., D.C., S.T., W.G., J.M.B., S.W., C.R.R., and J. L.

## Supporting Information

**SI Appendix, Fig. S1.**
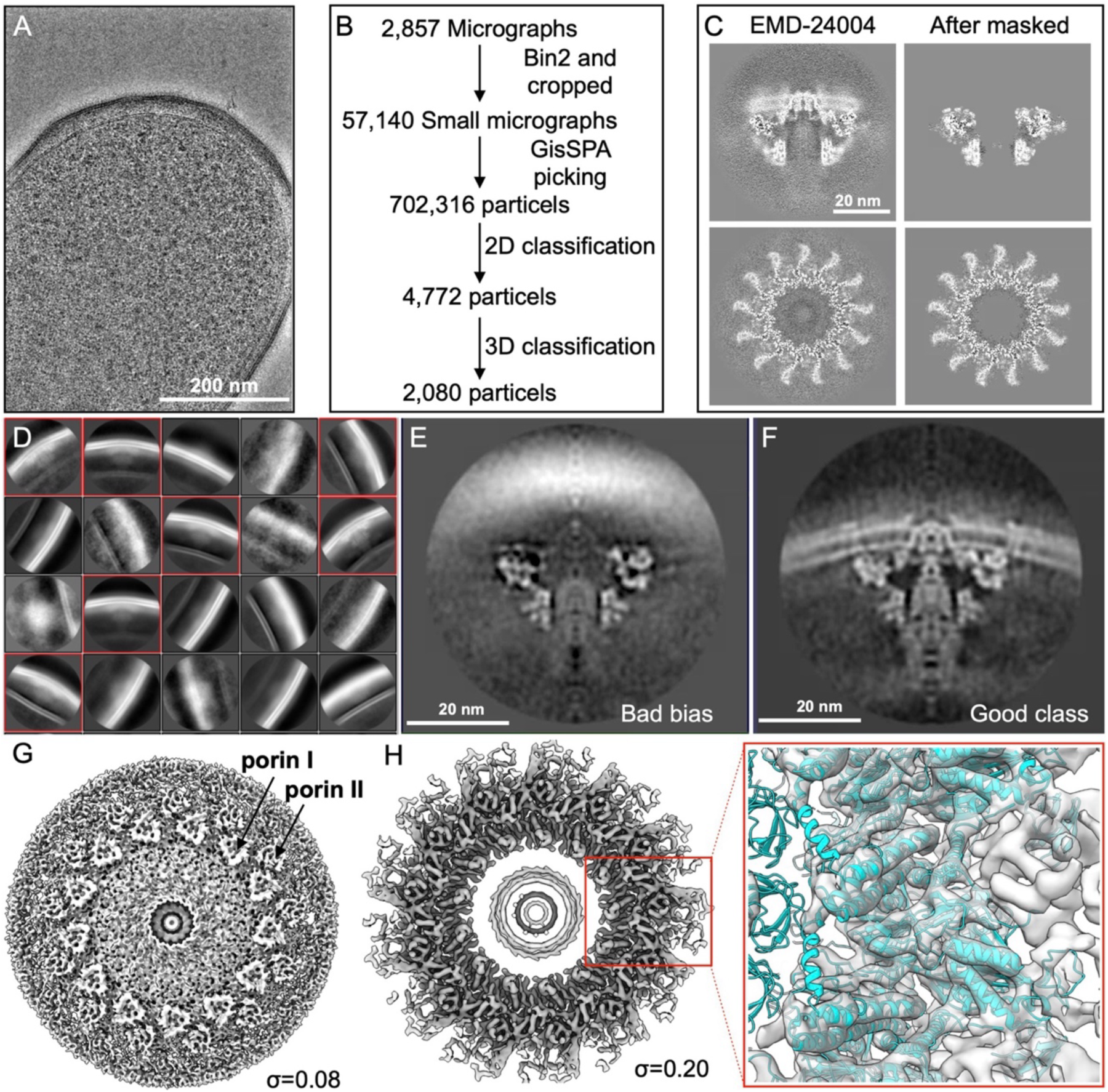
Initial cryo-EM analyses of Dot/Icm machines. (**A**) Representative micrograph acquired at the cell pole of *L. pneumophila*. (**B**) Workflow for particle selection by using GisSPA and RELION v3.1. A total of 2,857 raw micrographs were cropped into 1024×1024-pixel sub-micrographs and binned by a factor of 2 before being imported into GisSPA to optimize computational efficiency. Cryo-EM structure of the purified OMC (EMD-24004) was used as a high-resolution template, with detection thresholds set at a correlation coefficient of 6.5 and Euler angles sampled at 3° intervals. A total of 702,316 particles were detected, and their coordinates and orientation parameters were exported to RELION v3.1 for particle extraction. Subsequent 2D and 3D classification were performed to sort 2,080 high-quality particles. (**C**) Template used for GisSPA particle detection is based on EMD-24004 (left panel). The template was low-pass filtered to 8 Å and masked to remove low-resolution signals from detergent and dome regions (right panel). (**D**) Representative 2D classes. High-quality classes (red boxes) were selected for subsequent analyses. 3D classification shows a junk class (**E**) exhibiting only template features and a good class (**F**) displaying additional features. (**G**) Preliminary structure at a contour level of ∼0.08 σ. The porin structures in the outer membrane are indicated by arrows. (**H**) Surface view of the OMC structure at a contour level of ∼0.20 σ reveals secondary features. Fitting of the template model (PDB: 7MUD) into the density map was used to confirm the quality of the map.

**SI Appendix, Fig. S2.**
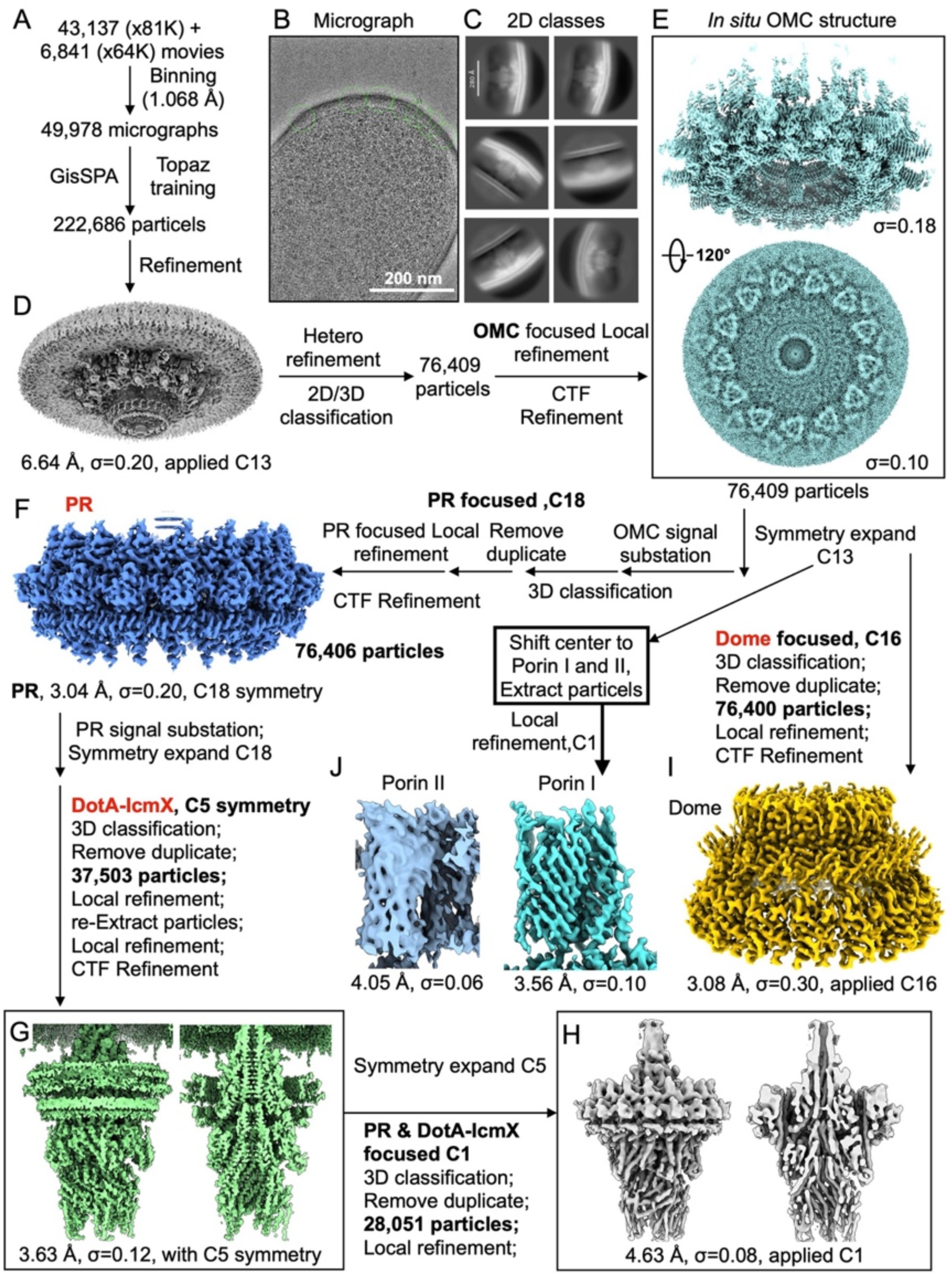
Overall image processing and structure determination of the Dot/Icm machine by *in-situ* cryo-EM. (**A**) Workflow for particle selection using a combination of GisSPA and Topaz. A total of 49,978 micrographs (43,137 at 81K magnification and 6,841 at 64K magnification) were binned to a final pixel size of 1.068 Å. Particles detected by GisSPA were used as a training set for Topaz based particle selection. (**B**) Representative particle selection results from iterative training and optimization with Topaz, enabling localization of 222,686 Dot/Icm particles at the cell pole (indicated by green circles). (**C**) Representative 2D classes of the extracted particles. (**D**) 3D reconstruction was generated through homogeneous refinement with C13 symmetry, using the preliminary structure in Figure S1G. Iterative heterogeneous refinement and 3D classification further optimized particle selection, yielding a high-quality subset of 76,409 particles. (**E**) The OMC structure with C13 symmetry at 2.96 Å resolution. (**F**) To resolve the PR, signal subtraction was applied to OMC-containing particles, followed by C13 symmetry expansion. Subsequent 3D classification identified PR features consistent with C18 symmetry. After removing duplicates, local refinement with C18 symmetry produced a 3.04 Å reconstruction. (**G**) A same strategy was applied to determine the structure of the DotA-IcmX complex. Of the initial particles, 37,503 (49%) exhibited clear channel density. These particles were refined with C5 symmetry to determine the *in-situ* structure at 3.63 Å resolution. (**H**) To examine the PR-Channel interactions, a symmetry-free reconstruction was performed. C5-expanded particles from the 3.63 Å dataset underwent 3D classification without symmetry constraints. The classes displaying both PR and channel features was refined with C1 symmetry, resulting in a 4.63 Å reconstruction. (**I**) Structural determination of the dome. OMC-expanded particles were subjected to 3D classification to identify features exhibited C16 symmetry. Final refinement with C16 symmetry produced a 3.08 Å structure. (**J**) Porin structure determination was conducted by focusing on Porins I and II. C13-expanded particles were re-centered on porin positions and refined locally with C1 symmetry to determine the structures.

**SI Appendix, Fig. S3.**
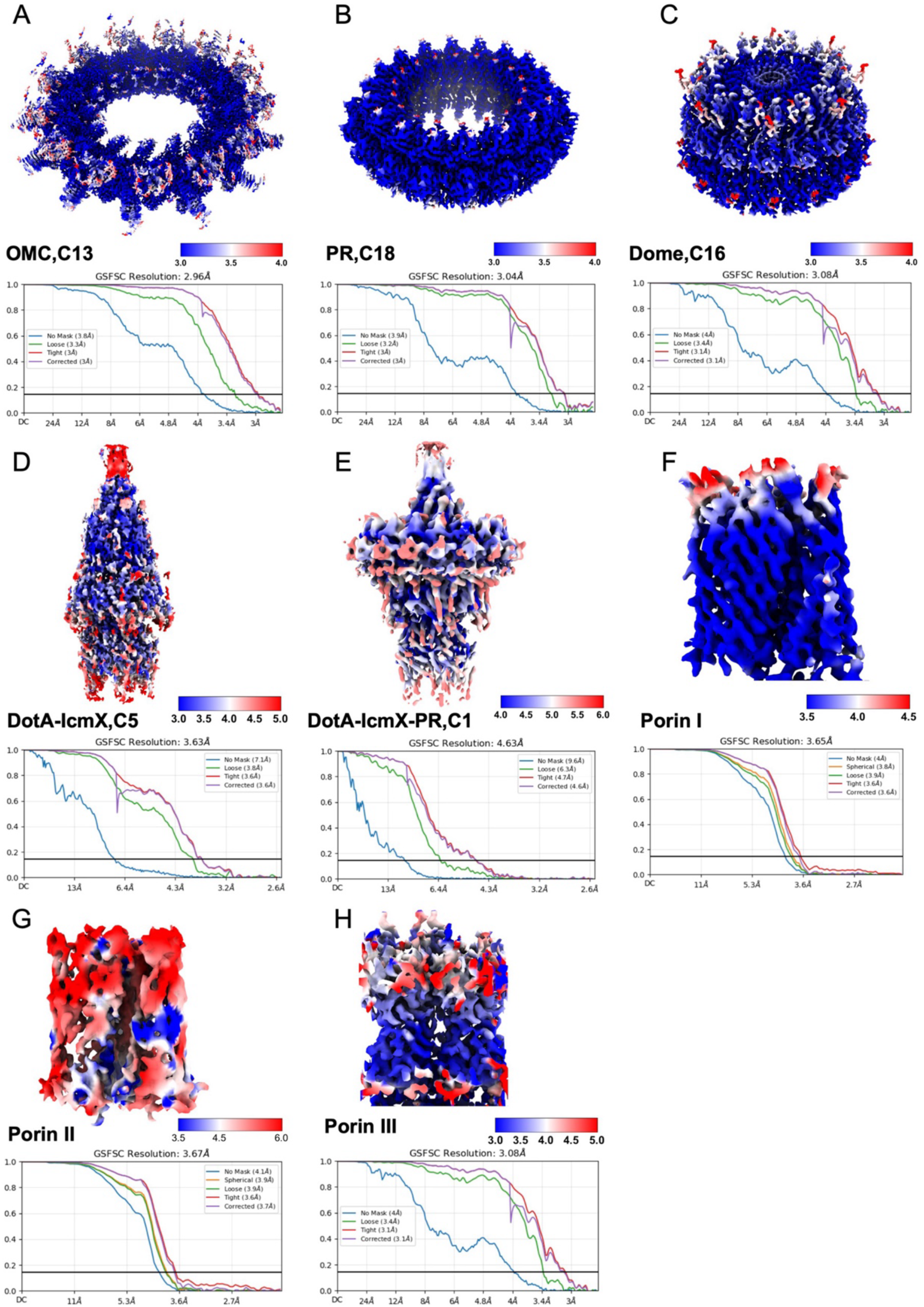
Resolution estimation of the Dot/Icm machine structure. Local resolution estimation maps and Fourier shell correlation (FSC) curves for each sub-complex: (**A**) OMC, (**B**) PR, (**C**) dome, (**D**) DotA-IcmX (C5), (**E**) DotA-IcmX-PR (C1), (**F**) porin I, (**G**) porin II, and (**H**) porin III.

**SI Appendix, Fig. S4.**
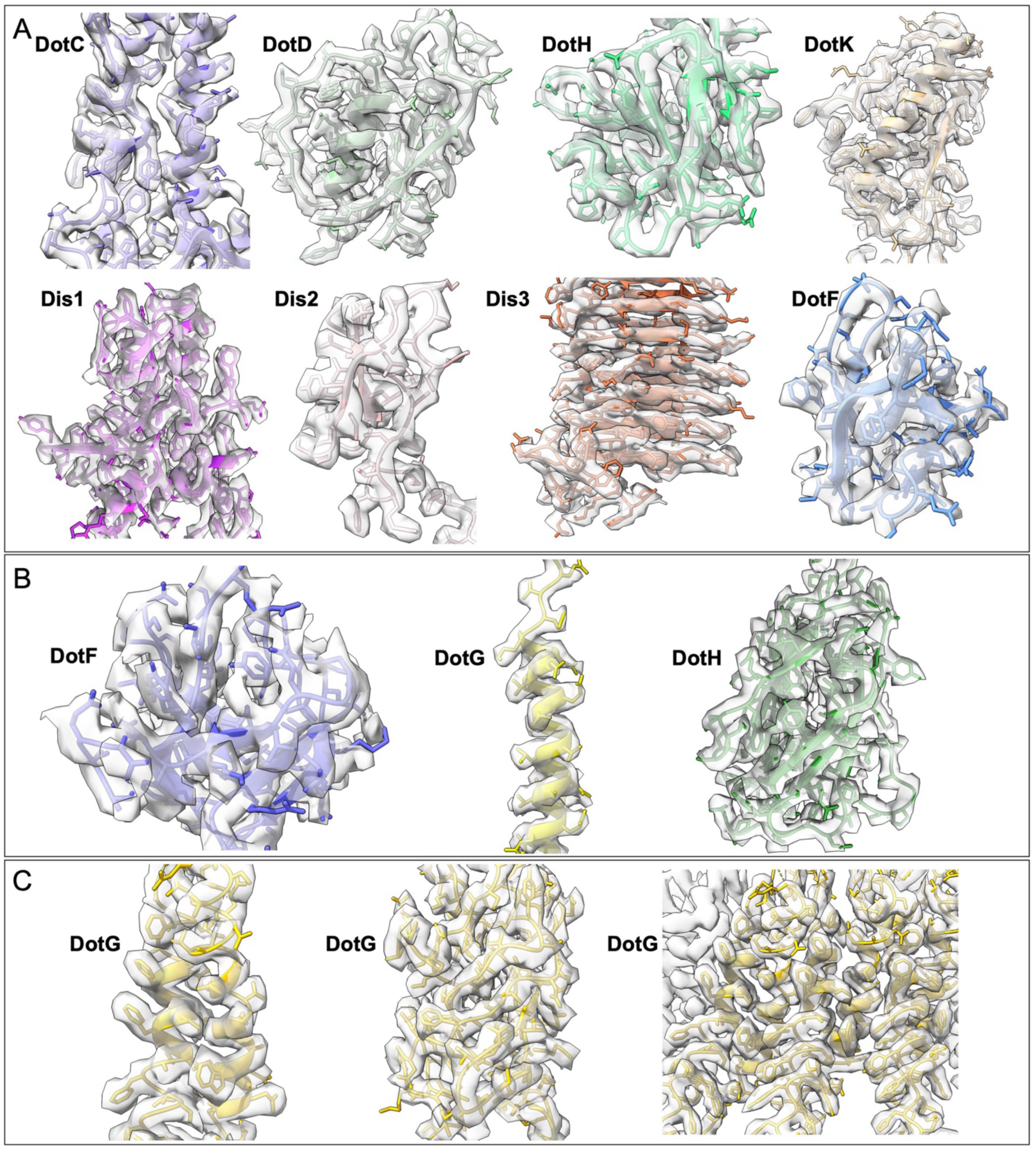
Model building of the OMC, PR, and dome. Maps displayed at a contour level of ∼0.10 σ in ChimeraX for each unit: (**A**) OMC, (**B**) PR, and (**C**) dome.

**SI Appendix, Fig. S5.**
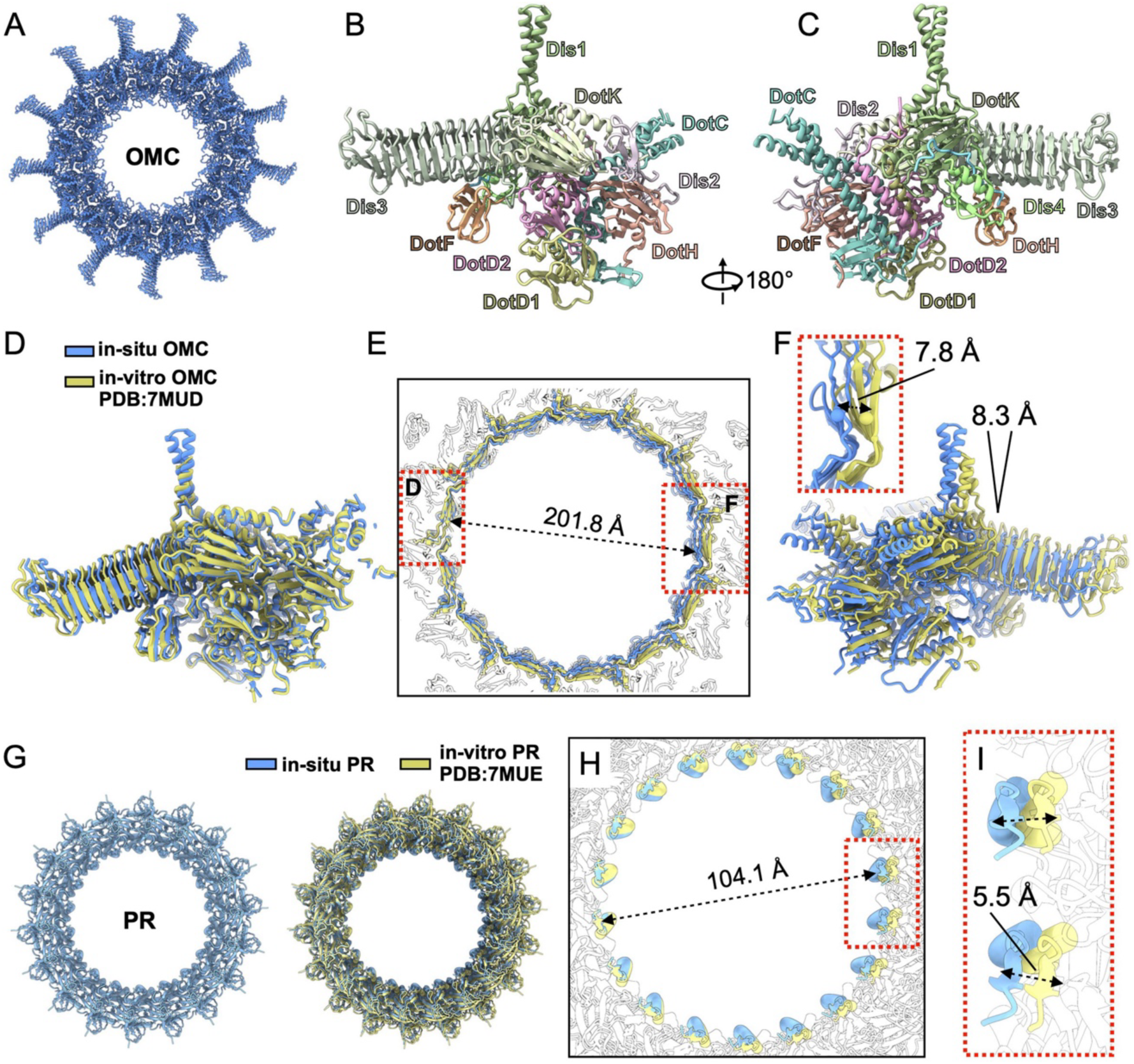
Structural comparison of the OMC and PR determined by *in-situ* and *in-vitro* approaches. (**A**) *In-situ* structure of the OMC is overall consistent with the cryo-EM structure of the OMC *in vitro*. (**B, C**) Two views of the asymmetric subunit of the OMC, with each component colored as indicated, respectively. (**D-F**) Structural comparison of the OMC determined by *in-situ* (blue) and *in-vitro* (yellow) methods reveals that the *in-situ* structure adopts a more compact conformation with a narrower inner diameter. Superposition of one asymmetric subunit (left side) of the OMC shows nearly perfect alignment (**D**), while the overall arrangement of asymmetric subunits displays noticeable conformational differences. The inner diameter of the OMC *in situ* is ∼7.8 Å narrower than that *in vitro* (**E**). (**F**) The asymmetric subunit on the right-side shifts by approximately 8.3 Å due to the overall conformational differences in the structure. (**G**) The overall architecture of the PR is conserved although a slight structural rearrangement is observed. Structural comparison of the PR determined by *in situ* and *in vitro* methods indicates that the *in-situ* structure also exhibits a more compact conformation (**H**). (**I**) The inner diameter of the PR *in situ* is ∼5.5 Å shorter than that *in vitro*.

**SI Appendix, Fig. S6.**
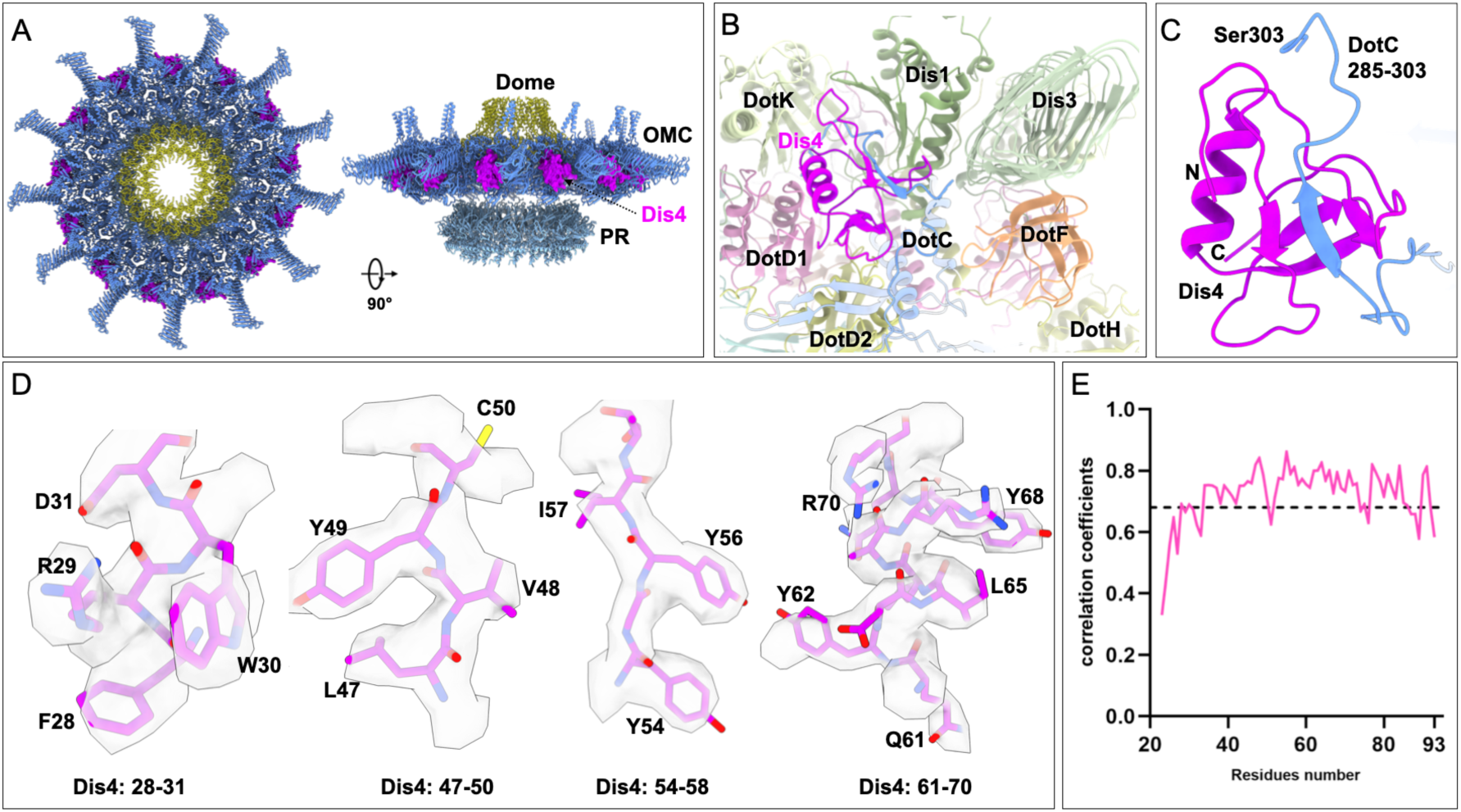
Identification of a new component Dis4 in the OMC. (**A**) Location of the new component, referred to as Dis4 (highlighted in purple), within the OMC, shown in two overall views. Dis4 is positioned at the center, between the spoke-like structures formed by Dis3. (**B**) Close-up view of Dis4 and its interactions with adjacent components. (**C**) Dis4 interacts with the C-terminus of DotC (residues 285–303), forming a three-stranded β-sheet that stabilizes its position within the OMC. (**D**) Model building of Dis4 at a contour level of ∼0.10 σ in ChimeraX. (**E**) Analysis of correlation coefficients per residue for Dis4, showing the quality of the model fitting.

**SI Appendix, Fig. S7.**
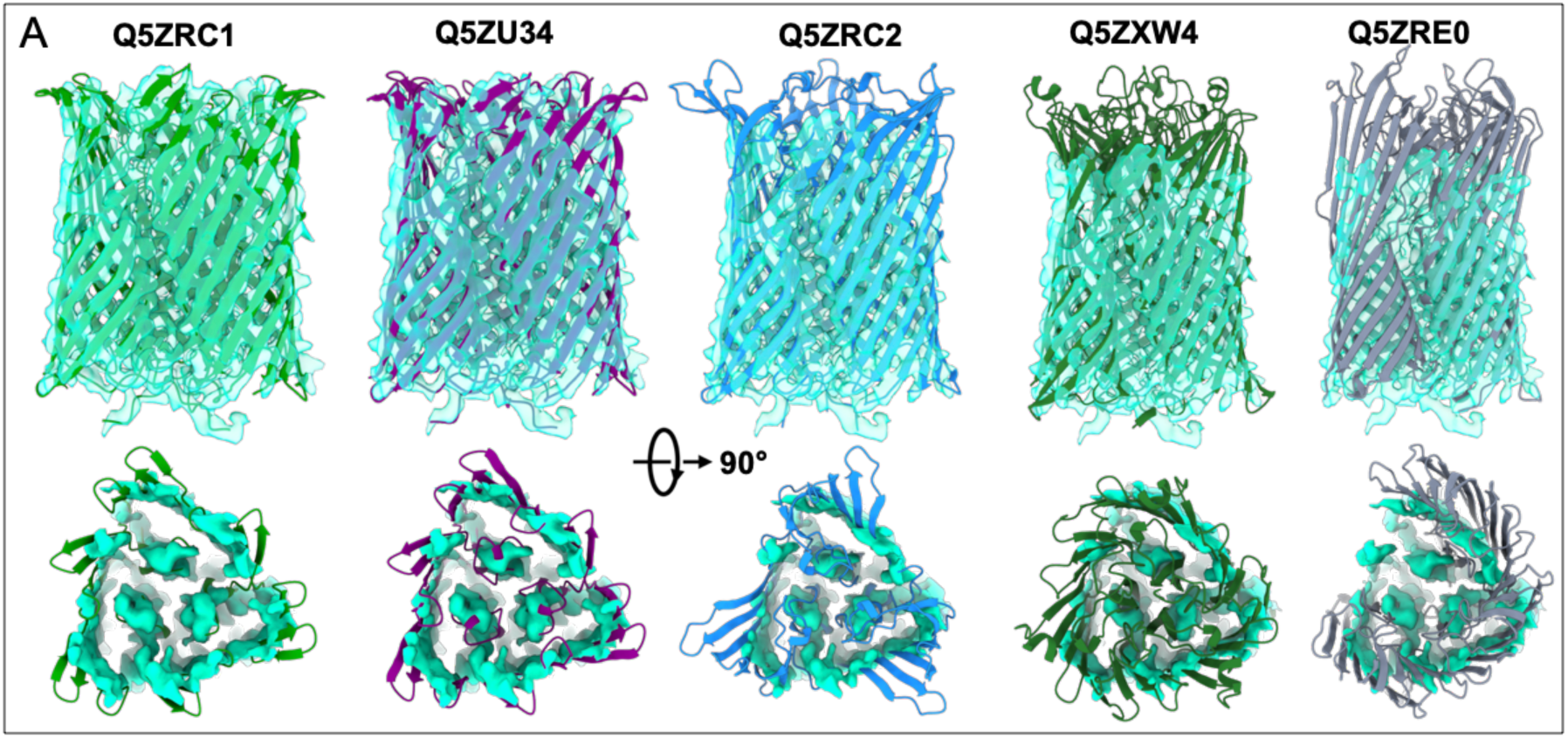
Identification of porin proteins. For porin sequence identification and structural modeling, we screened all β-barrel-like proteins in *L. pneumophila* (Taxon ID: 272624). Predicted structures by AlphaFold3 were fitted into the density map individually. Porin I, a five 10- stranded β-barrel protein, matched the density map features. The trimeric structure of the MOMP (UniProt: Q5ZRC1; gene locus: *lpg2961*) exhibited the best fit.

**SI Appendix, Fig. S8.**
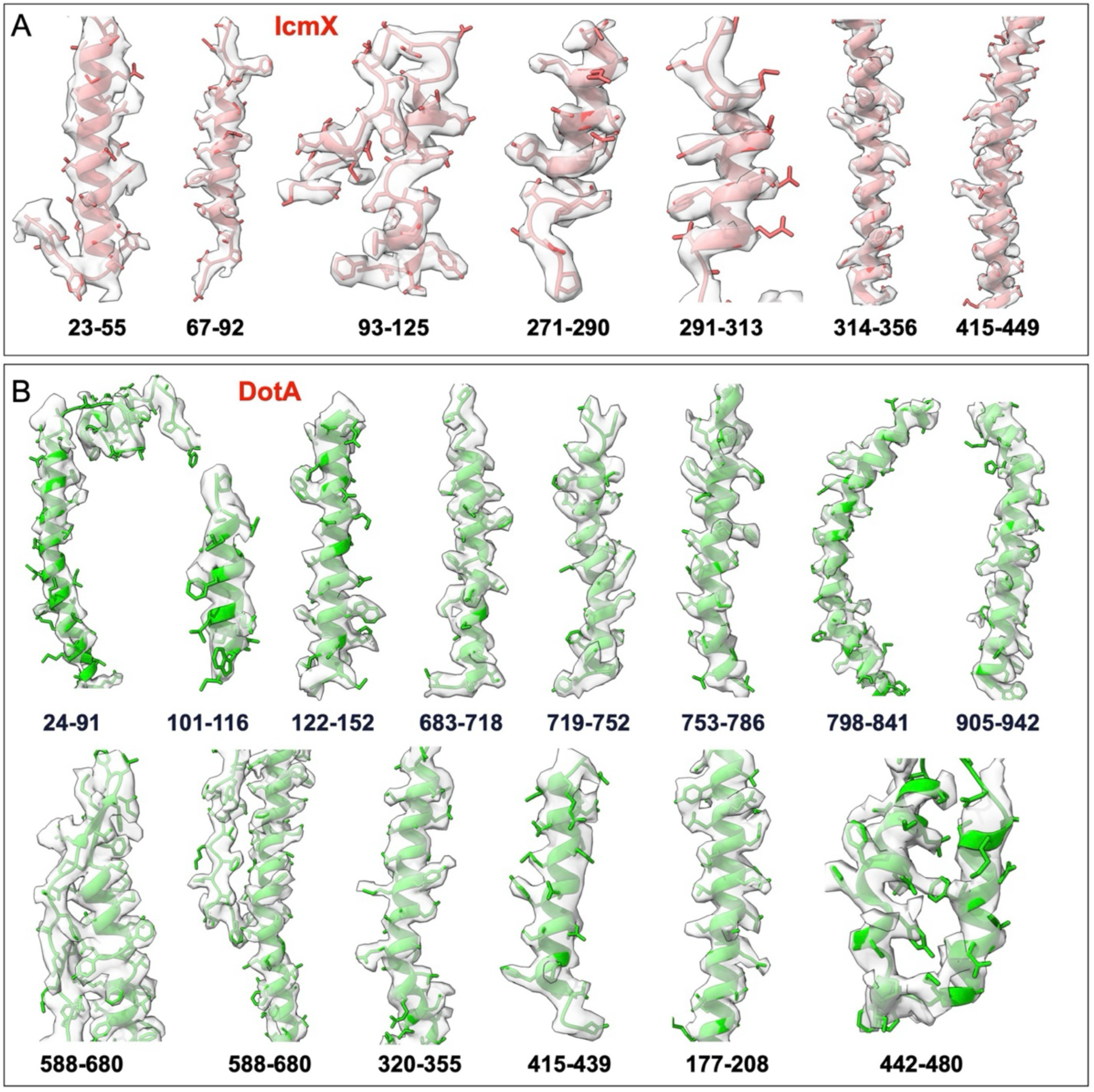
Model building of DotA and IcmX. Maps displayed at a contour level of ∼0.10 σ in ChimeraX for IcmX (**A**) and DotA (**B**). The corresponding residue regions are indicated.

**SI Appendix, Fig. S9.**
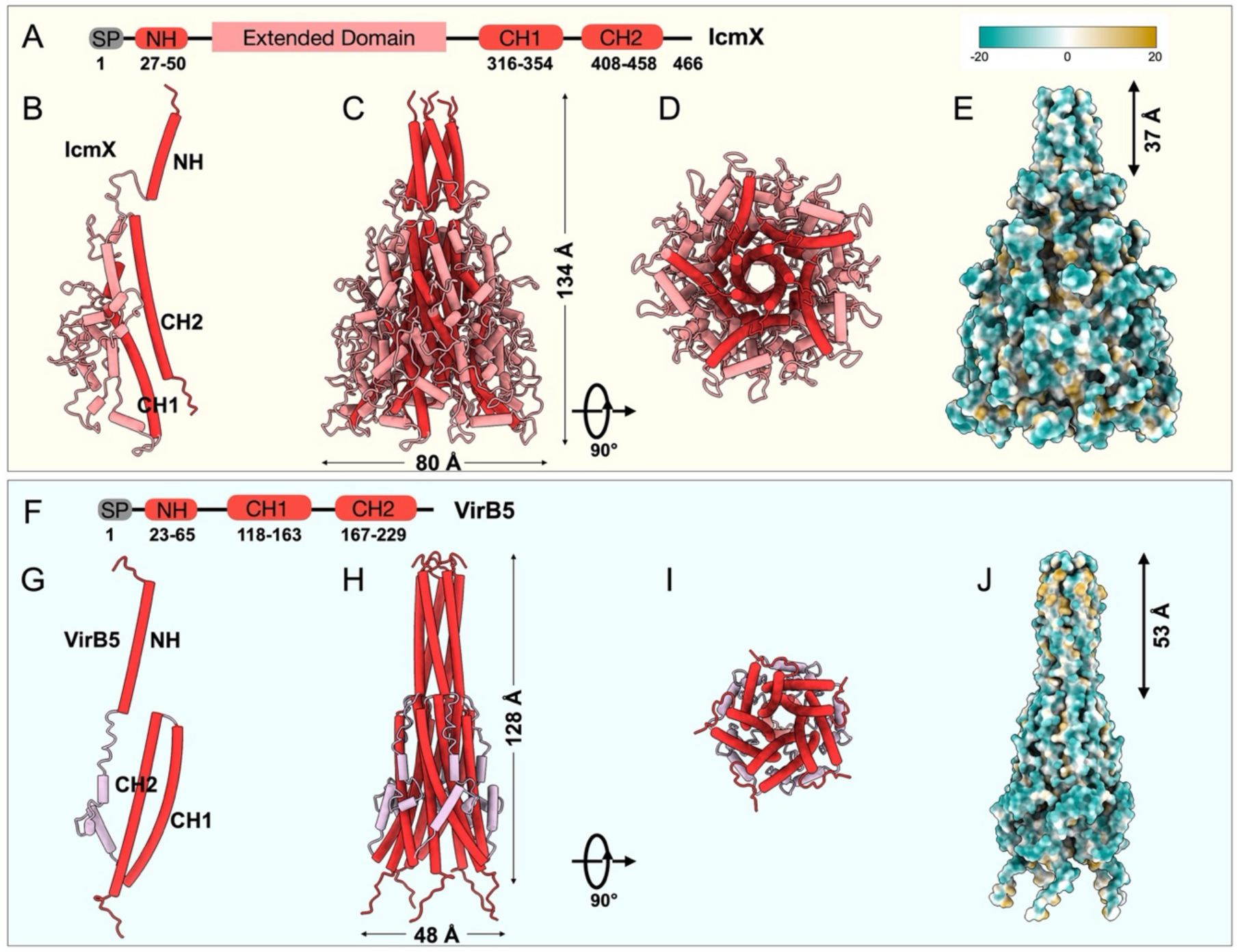
Structural comparison of IcmX and VirB5. (**A**) Schematic diagram of IcmX domain structure. IcmX includes a signal peptide (SP) and conserved N-terminal (NH) and C-terminal (CH1 and CH2) α-helices, which share structural similarity with VirB5 (panels **F** and **G**), a conjugation T4SS component. IcmX features a larger extended domain compared to VirB5. (**B**) Monomeric structure of IcmX. Conserved NH, CH1, and CH2 helices are colored red, while the extended domain is highlighted in light red. (**C, D**) Pentameric structures of IcmX. Side view (**C**) and top view (**D**) are presented as cylinder diagrams. The extended domain makes IcmX (∼80 Å) much wider than VirB5 (∼48 Å). (**E**) Surface properties of IcmX shown as lipophilicity potential. (**F**) Schematic diagram of VirB5 domain organization. VirB5 comprises a signal peptide (SP) and three primary α-helices (NH, CH1, and CH2) at its termini. (**G**) Monomeric structure of VirB5. Shown as a ribbon diagram, with structural homologs of IcmX’s α-helices colored red. (**H**, **I**) Pentameric structures of VirB5. Side view (**H**) and top view (**I**) are displayed as cylinder diagrams to illustrate its assembly. (**J**) Surface properties of VirB5. Hydrophobic and hydrophilic surface properties are shown for comparison.

**SI Appendix, Fig. S10.**
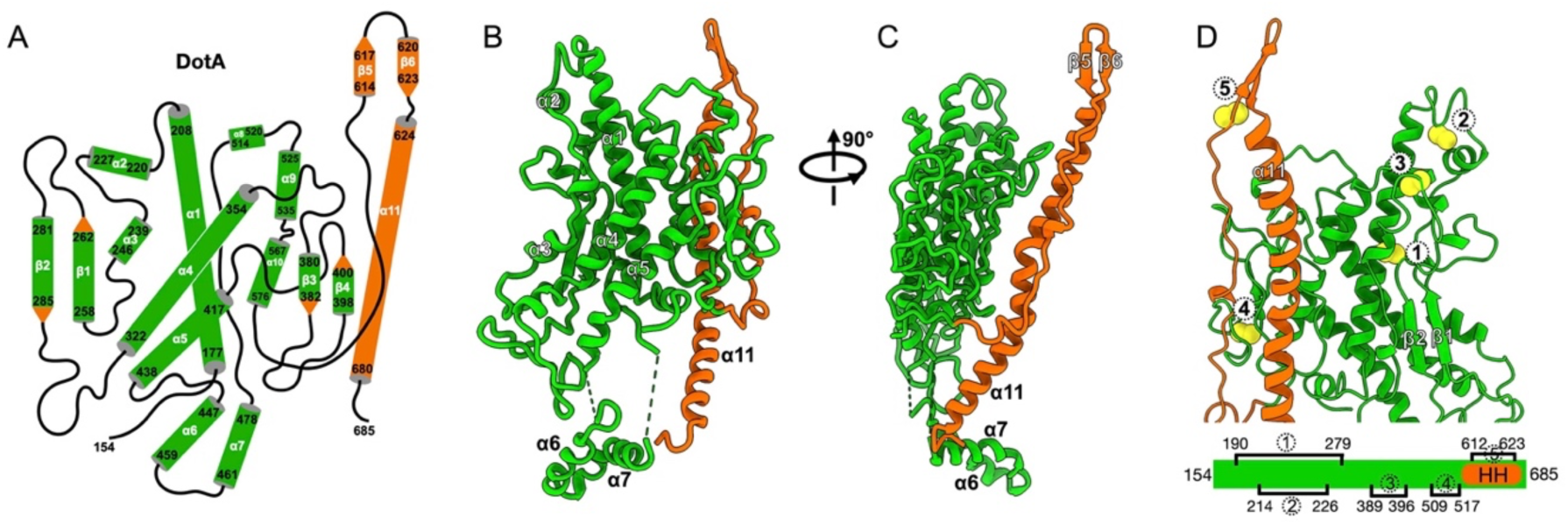
Structural details of the DotA extended domain. (**A**) Structural topology of the DotA extended domain. (**B, C**) Two views of the DotA extended domain, respectively. (**D**) The disulfide bonds in the DotA extended domain are indicated by yellow balls.

**SI Appendix, Fig. S11.**
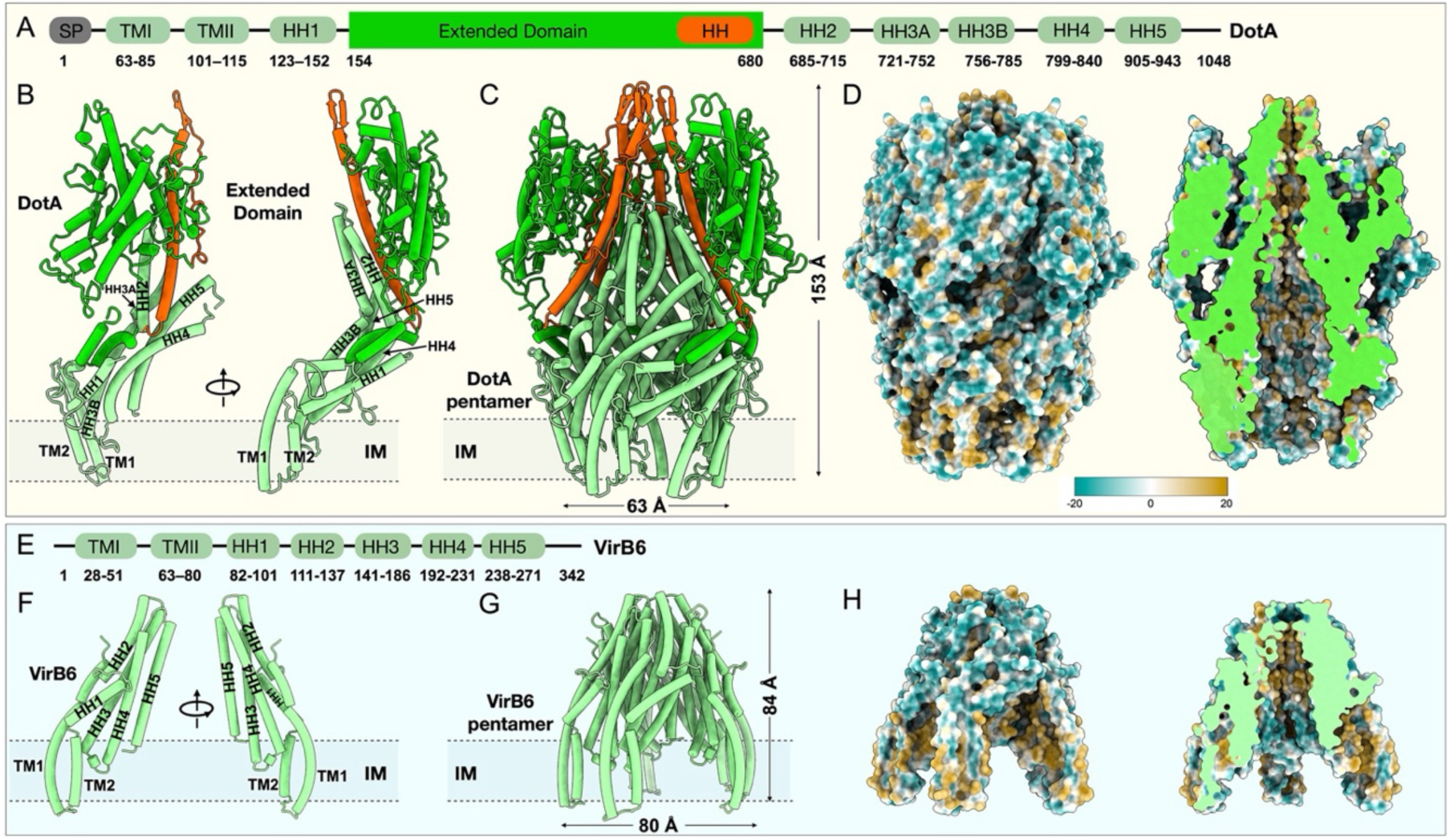
Structural comparison of DotA and VirB6. (**A**) Schematic diagram of DotA domain organization. DotA includes a signal peptide (SP), two N-terminal transmembrane α- helices (TMI and TMII), and five long hydrophobic helices (HH1–HH5). These features exhibit structural similarity to VirB6, conjugation T4SS component (see panels **E** and **F**). (**B**) Monomeric structure of DotA. The transmembrane helices (TMI, TMII) and hydrophobic helices (HH1–HH5) are shown in light green. The extended domain is highlighted in green, with one hydrophobic helix and a β-hairpin shown in orange. (**C**) Pentameric structure of DotA, depicting its assembly. One of the front extended domains is hidden to expose the internal architecture of the assembly. (**D**) Surface properties of DotA. The left panel shows an overall view, while the right panel presents a cross-sectional view along the central axis. (**E**) Schematic diagram of VirB6 domain organization. VirB6 features two N-terminal transmembrane α-helices (TMI and TMII) and five hydrophobic helices (HH1–HH5) at the C-terminal, which are structurally similar to DotA, but helixes length is shorter than virB6. (**F**) Monomeric structure of VirB6. The transmembrane helices (TMI, TMII) and hydrophobic helices (HH1–HH5) are colored light green. (**G**) Pentameric structure of VirB6. Shown as a comparison to DotA (**C** and **D**), revealing that DotA is larger, wider, and more enclosed compared to the structure of VirB6. (**H**) Surface properties of VirB6. The left panel illustrates an overall view, while the right panel shows a cross-sectional view of the central axis.

**SI Appendix, Fig. S12.**
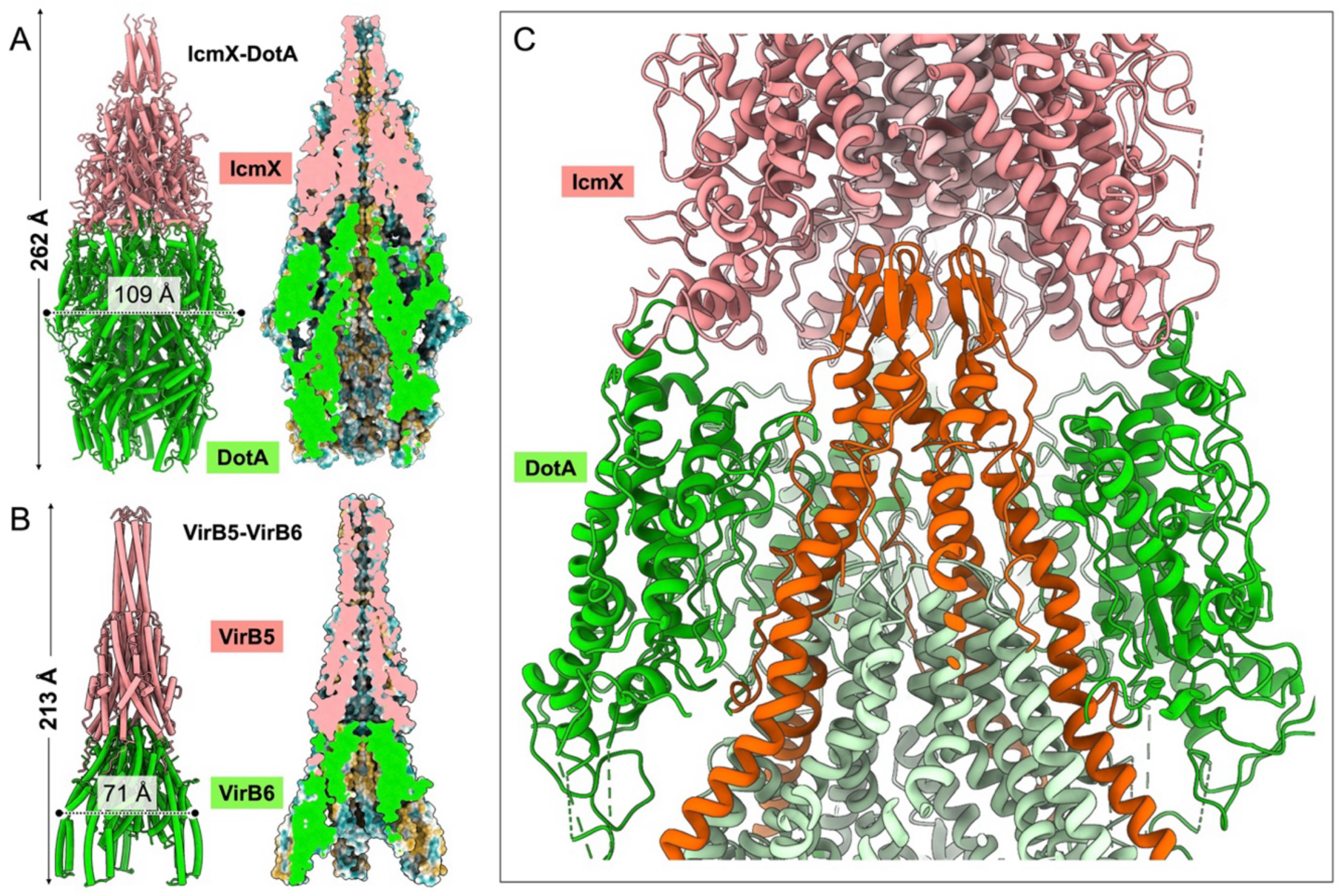
Structural comparison of DotA-IcmX and VirB5-VirB6. (**A**) Structure of the DotA-IcmX complex. IcmX and DotA are shown in red and green, respectively. The left panel presents the overall view as a cartoon model, while the right panel provides a cross-sectional view through the center of the complex. The height and width of the assembly are labeled. (**B**) Structure of the VirB5-VirB6 complex. VirB5 and VirB6 are shown in red and green, respectively. A cartoon model is shown in left panel and a cross-sectional view through the VirB5-VirB6 complex in right panel, respectively. The height and width of this assembly are labeled for comparison with IcmX and DotA. (**C**) Detailed view of the DotA-IcmX interface. An enlarged view emphasizes the extensive interaction between their extended domains. The extended domains of DotA and IcmX are highlighted in green and orange, respectively, to underscore their critical role in the interaction.

**SI Appendix, Fig. S13.**
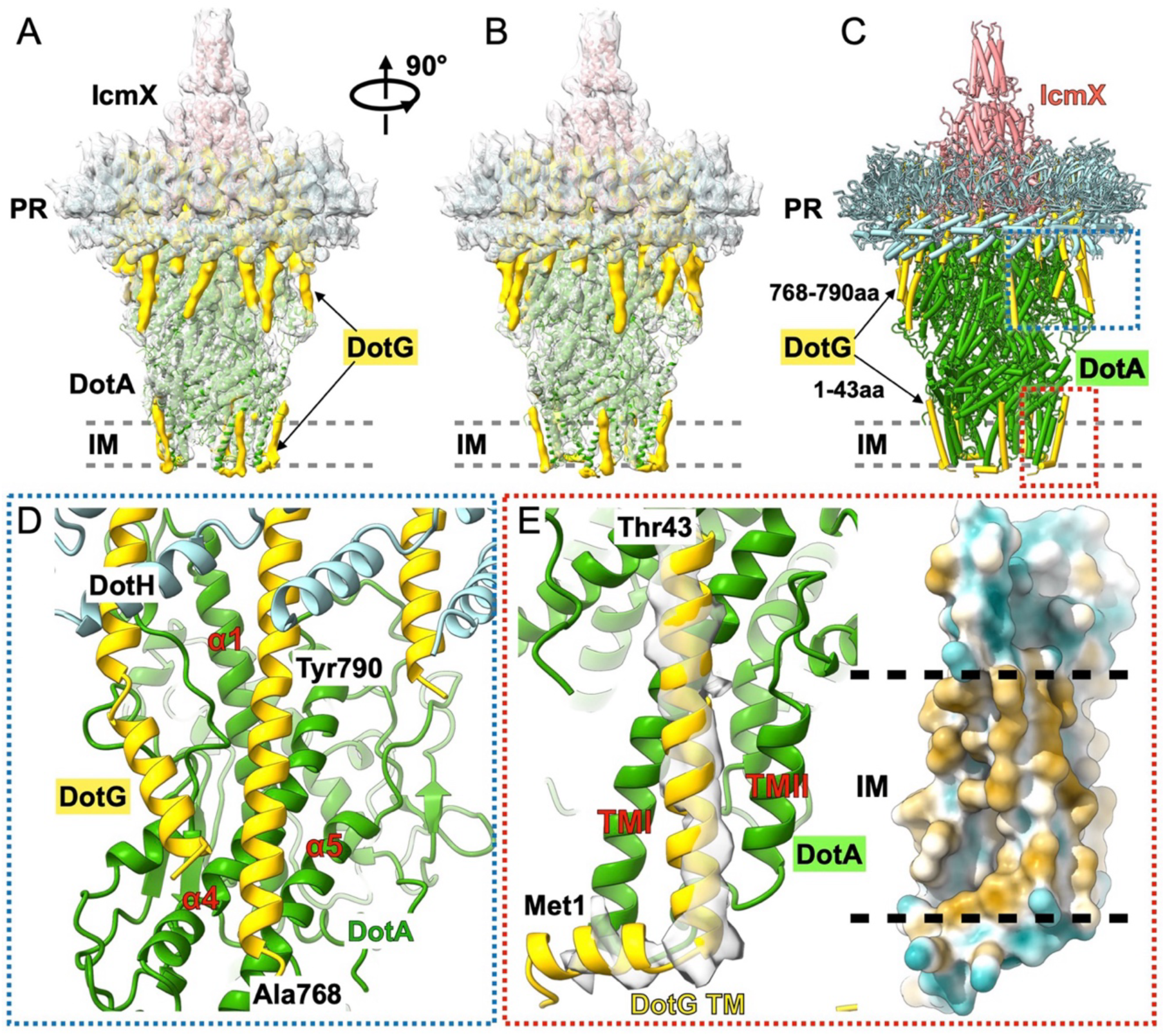
Interactions between DotA and DotG. (**A, B**) Two views of the density map of PR and DotA-IcmX with C1 symmetry. Except for the model of DotA-IcmX and PR, there is an extra density corresponding to DotG (yellow). Eighteen copies of DotG (residues 791-824) adopt C18 symmetry that constitute the interior of PR. The extra density protruding from PR and attached on the DotA extended domain surface displays no symmetry property. In addition, five copies of helix-like density surround the DotA transmembrane helix. (**C**) The model of DotG. Due to resolution limitations, the structural modeling was based on AlphaFold predictions and adjusted by density. (**D, E**) Close-up views of the DotA and DotG interactions. (**D**) The DotG helices (residues 768-790) protruding from PR are attached to the surface of the DotA extended domain. (**E**) A close-up view of the interaction between DotA (TMI and TMII) and the transmembrane region of DotG (residues 1-43).

**SI Appendix, Fig. S14.**
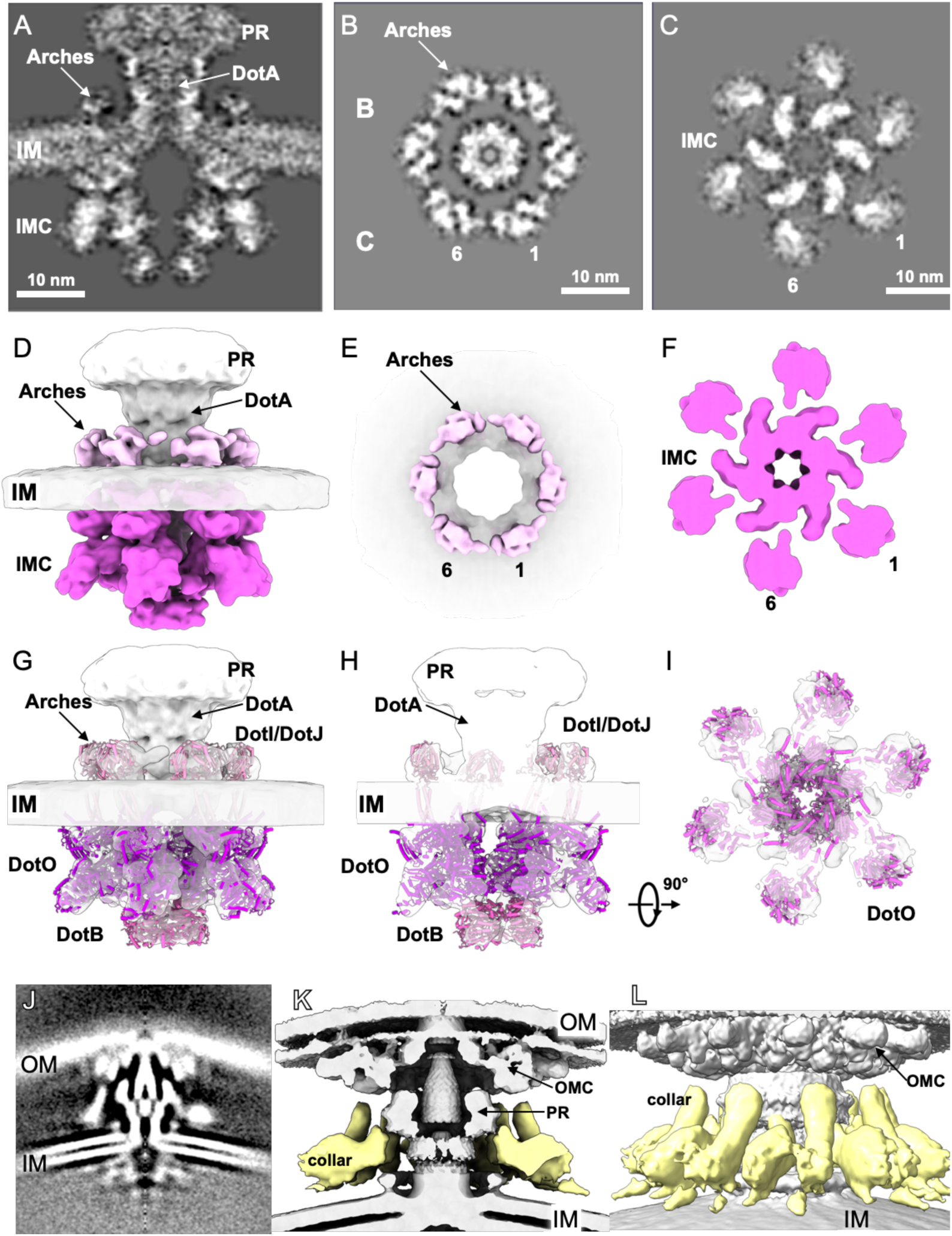
*In-situ* cryo-ET structure of the inner membrane complex (IMC), arches, and collar of the Dot/Icm machine. (A) A central section of the cryo-ET structure of the IMC. Sub-tomogram averaging and 3D classification revealed a C6 symmetric structure on the cytoplasmic side and an arches-like structure on the periplasmic side. Top view of the arches-like structure (B) and the IMC structure (C). (D) 3D rendering of the cryo-ET density. The IMC is highlighted in magenta. (E, F) Top view of the 3D rendering density of the arches (E) and IMC (F). (G-I) Structural modeling of the IMC and arches. A dimer of hexamers for DotO was fitted into the IMC, and a hexamer of DotB was fitted at the bottom. The arches were modeled based on previous reports, consisting of DotI-DotJ. (J) A central section of our cryo-ET structure showing the collar-like density (yellow), which exhibits two distinct forms: one likely connecting the OMC and IM, and the other, positioned slightly lower, connecting the PR and extending toward the IM. (K, L) Two view showing these two forms of densities as the 3D surface.

**SI Appendix, Fig. S15.**
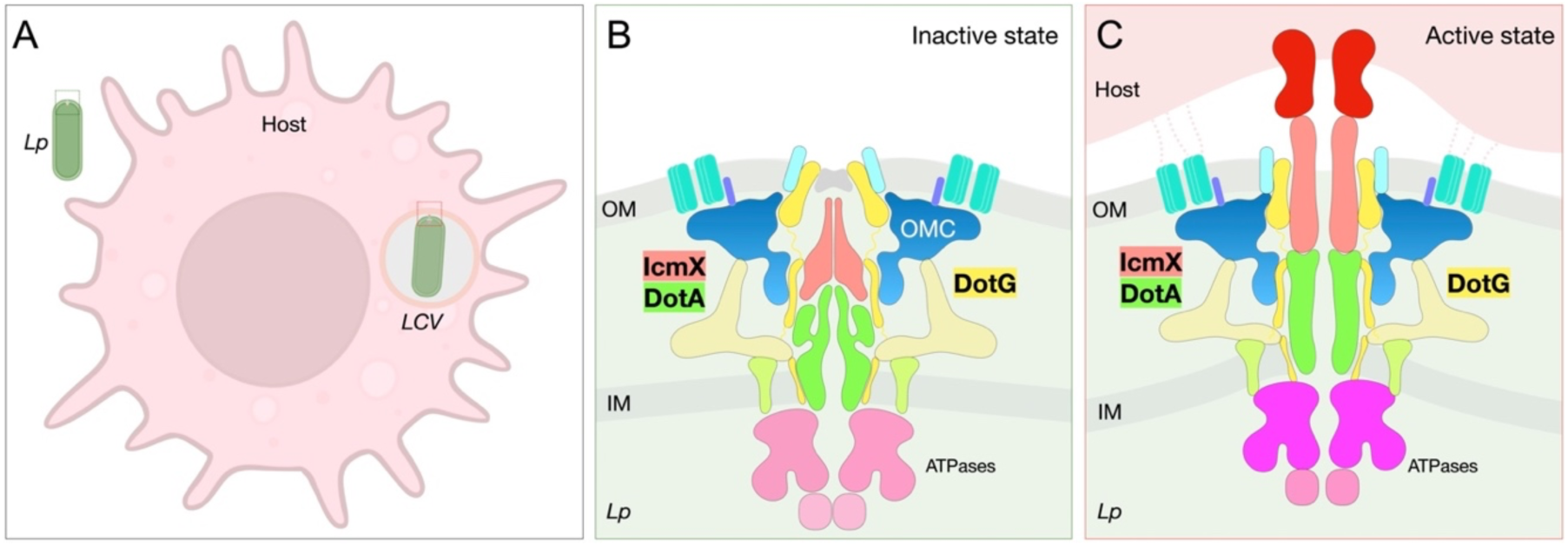
Model of the Dot/Icm T4SS activation upon host contact. (**A**) Interactions between *L. pneumophila* and macrophage. One *L. pneumophila* cell is located outside of the macrophage. Another one resides in the LCV. (**B**) Dot/Icm machine structure in an inactive state. (**C**) Host contract triggers extensive conformational changes in the Dot/Icm machine to form a transenvelope conduit, enabling direct effector protein translocation from the bacterial cytosol to recipient cell.

**SI Appendix, Table S1.**
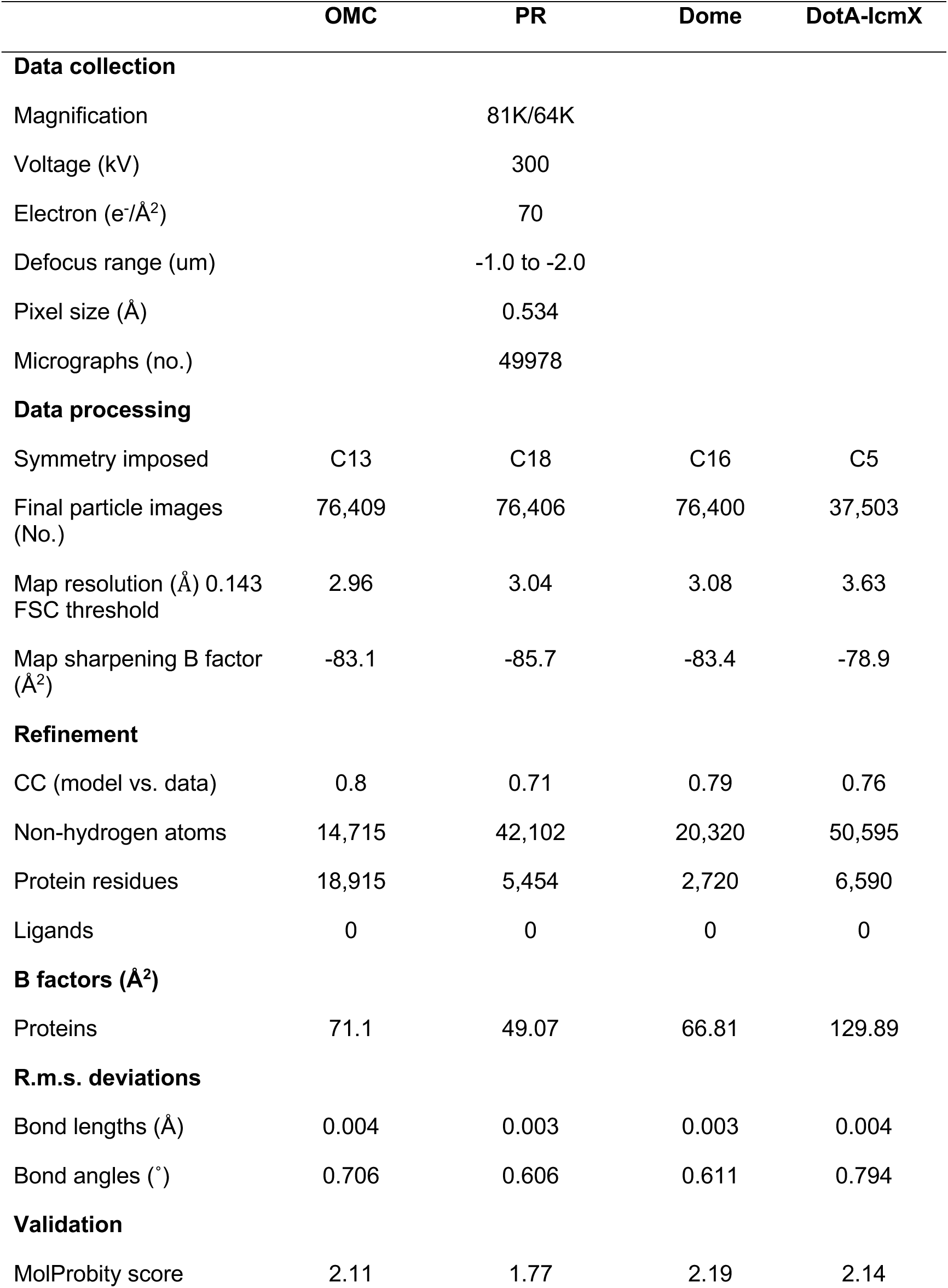
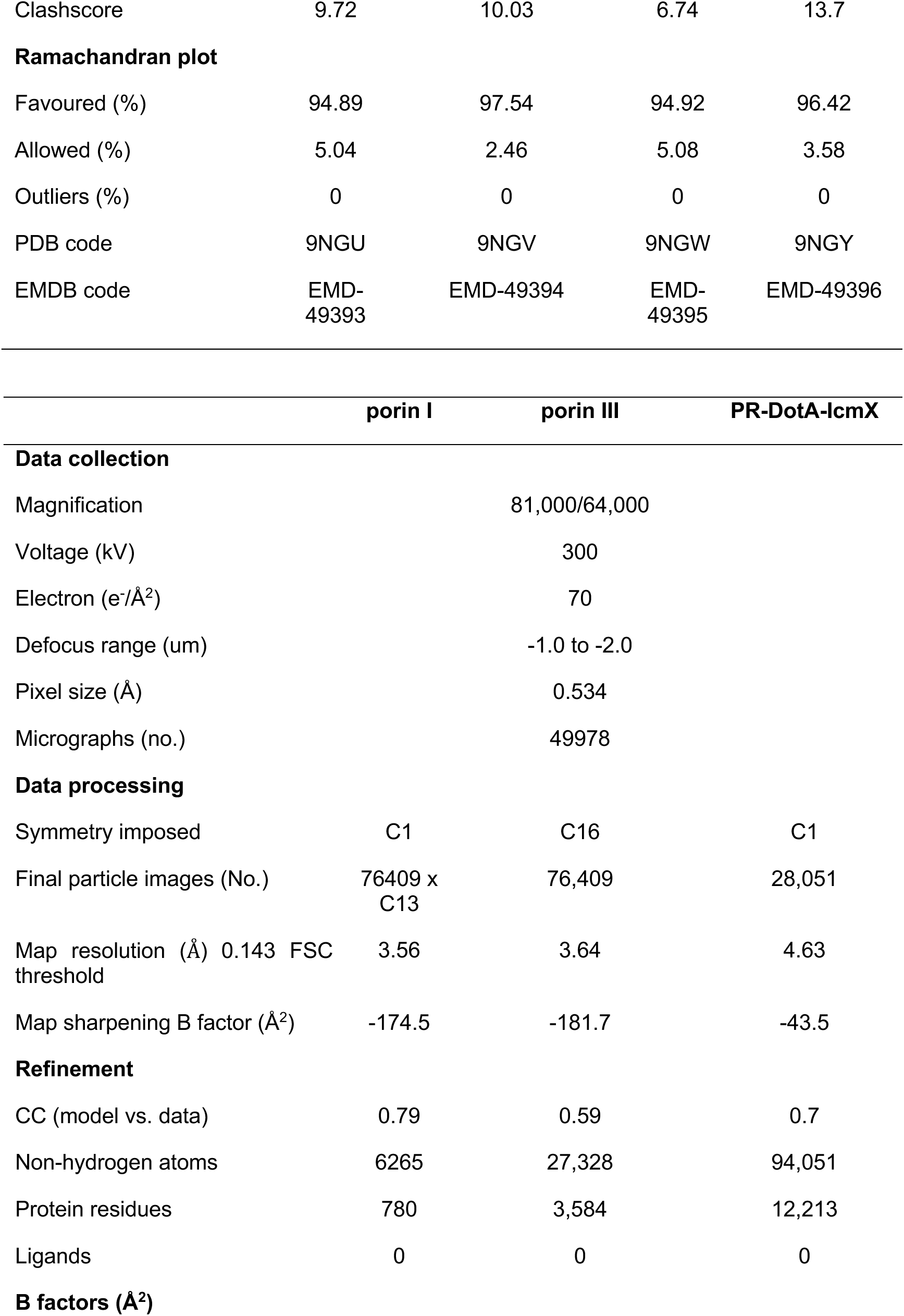
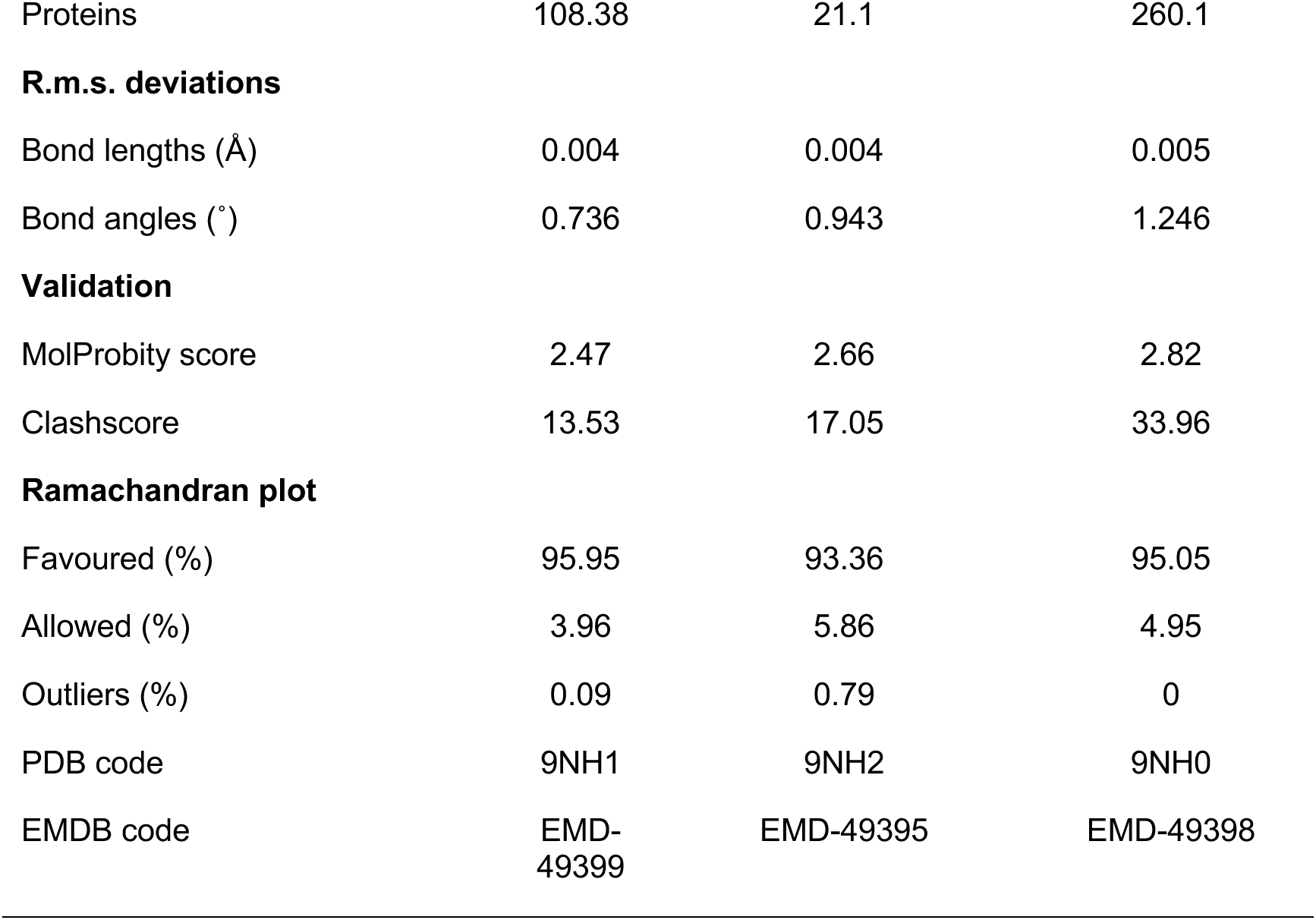
Data collection and structure refinement statistics

**SI Appendix, Table S2.** Statistics of residues modeled in the Dot/Icm machine

| Protein | Length | Uniport | OMC | PR | Dome | Protochannel |
| --- | --- | --- | --- | --- | --- | --- |
| DotG | 1048 | Q5ZYC1 |  | 767-824 | 862-1048 | 2-43 |
| DotH | 361 | Q5ZYC2 | 271-361 | 104-263 |  |  |
| DotF | 269 | Q5ZYC0 | 208-266 |  |  |  |
| DotD | 163 | Q5ZS45 | 24-162 |  |  |  |
| DotC | 303 | Q5ZS44 | 26-39, 53-161, 170-303 |  |  |  |
| DotK | 189 | Q5ZYC5 | 40-188 |  |  |  |
| Dis1 | 249 | Q5ZXS4 | 42-235 |  |  |  |
| Dis2 | 124 | Q5ZXA9 | 40-115 |  |  |  |
| Dis3 | 320 | Q5ZRN3 | 29-320 |  |  |  |
| **Dis4** | 93 | A0AAN5PM39 | 22-93 |  |  |  |
| **Porin I** | 288 | Q5ZRC1 | 23-288 |  |  |  |
| **Porin II** | 288 | Q5ZRC1 | 23-288 |  |  |  |
| **Porin III** | 245 | ---- |  |  |  |  |
| **lcmX** | 466 | Q5ZS30 |  |  |  | 23-148, 168-458 |
| DotA | 1048 | Q5ZS33 |  |  |  | 24-154, 160-439, 442-480, 483-845, 901-945 |

**SI Appendix, Table S3.**
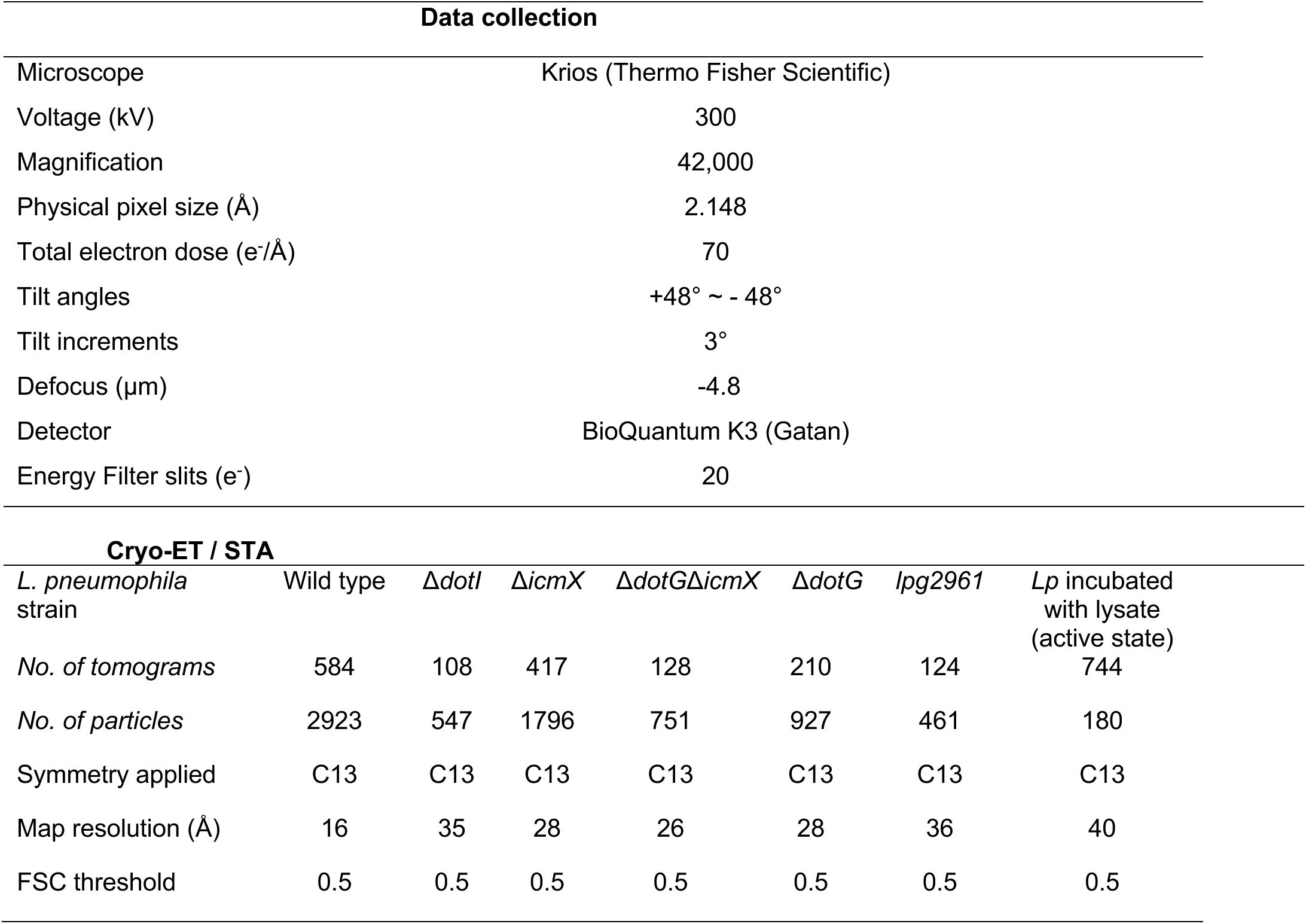
Cryo-ET data collection and structure refinement statistics

***SI Appendix,* Movie S1.** An animation illustrates the near-atomic structure of the Dot/Icm machine in *L. pneumophila*.

***SI Appendix,* Movie S2.** Tomogram of a FIB-milled lamella from *L. pneumophila* inside a macrophage shows a bacterium tethered to the LCV membrane via a Dot/Icm machine. The movie highlights a Dot/Icm machine forming a conduit, seen as additional density between the bacterial outer membrane and the LCV membrane, suggesting a pathway for effector protein translocation. Hcyt: host cytoplasm, Mt: host mitochondria.

***SI Appendix,* Movie S3.** Side-by-side tomograms of *L. pneumophila* cell envelopes illustrate the structural differences between activated and inactivated Dot/Icm machines. The left tomogram shows multiple activated Dot/Icm machines embedded within the bacterial cell envelope, each forming an extracellular density extending beyond the outer membrane. In contrast, the right tomogram shows inactivated Dot/Icm machines, which lack the extracellular density, representing a non-secreting state.

***SI Appendix,* Movie S4.** An animation illustrates an activation model of the Dot/Icm machine through host contact.

